# SCLC TumorMiner: A Genomics Platform for Small Cell Lung Cancer Precision Oncology

**DOI:** 10.1101/2025.07.07.663501

**Authors:** Fathi Elloumi, Anjali Dhall, Daiki Taniyama, Augustin Luna, Yasuhiro Arakawa, Sudhir Varma, Yanghsin Wang, Anisha Tehim, Mark Raffeld, Kenneth Aldape, Christophe Redon, Roshan Shresta, William Reinhold, Mirit Aladjem, Jaydira Del Rivero, Nitin Roper, Anish Thomas, Yves Pommier

**Author notes:** Equal contributions (Fathi Elloumi, Anjali Dhall).

## Abstract

Small cell lung cancer (SCLC) is among the most aggressive malignancies. Unlike many other cancers, it is not represented in The Cancer Genome Atlas, and available datasets are fragmented across institutions, disease stages, and treatment settings. RNA sequencing provides a powerful and cost-effective approach, but the high dimensionality of transcriptomic data and the heterogeneity of patient cohorts pose significant challenges. To address such challenges, we developed SCLC TumorMiner (https://discover.nci.nih.gov/SclcTumorMinerCDB/), which includes 50 tumor samples from relapse patients at the National Cancer Institute (NCI) and 154 samples from untreated patients at the University of Cologne and Tongji University. SCLC TumorMiner enables molecular classification, genomic pathway analyses, risk stratification, identification of predictive cell-surface biomarkers such as DLL3 or TROP2, and drug-response biomarkers such as SLFN11. SCLC TumorMiner illustrates profound differences between untreated and relapse patient samples. Additionally, “MyPatient”, one of SCLC TumorMiner’s modules, is presented as a medical assistant application prototype.

## INTRODUCTION

Small cell lung cancers are rapidly proliferative tumors often diagnosed with metastases (Extensive-stage; ES-SCLC) ^1^. They account for approximately 13%–15% of all lung cancer cases and remain one of the most lethal malignancies. Survival is less than one year for most patients, and 95% of patients die within 5 years despite initial response to cisplatin-etoposide combination therapy. Upon relapse, tumors evolve and become resistant to chemotherapy, reflecting profound changes in their biology and genomic network.

In addition to mutations of the tumor suppressors TP53 and RB1 in 80% of SCLC, selective activation of transcription factors characterizes the disease ^2,3^. At diagnosis, SCLCs have been divided into two groups. 70-80% are neuroendocrine (NE), driven by two pioneer neurogenic transcription factors, ASCL1 (Acheate-scute homolog 1) and NEUROD1 (Neurogenic differentiation factor 1). The remaining non-NE SCLCs are driven by two other transcription factors POU2F3 (POU class 2 homeobox 3) and YAP1 (Yes-associated protein 1). Consequently, SCLCs have been classified into 4 subsets: SCLC-A, SCLC-N, SCLC-P, and SCLC-Y ^3^ (NAPY classification ^3,4^). Additionally, an Inflamed/Immune subgroup has been identified as SCLC-I ^5^. These classifications are well supported in untreated tumors and SCLC cell lines ^3,4^ (see Figure 1B), but are not routinely applied in clinical practice, where diagnosis and classification rely primarily on neuroendocrine marker immunohistochemistry for synaptophysin (encoded by the *SYP* gene), chromogranin (encoded by *CHGA*), and neural cell adhesion molecule (encoded by *NCAM1*).

**Figure 1.**
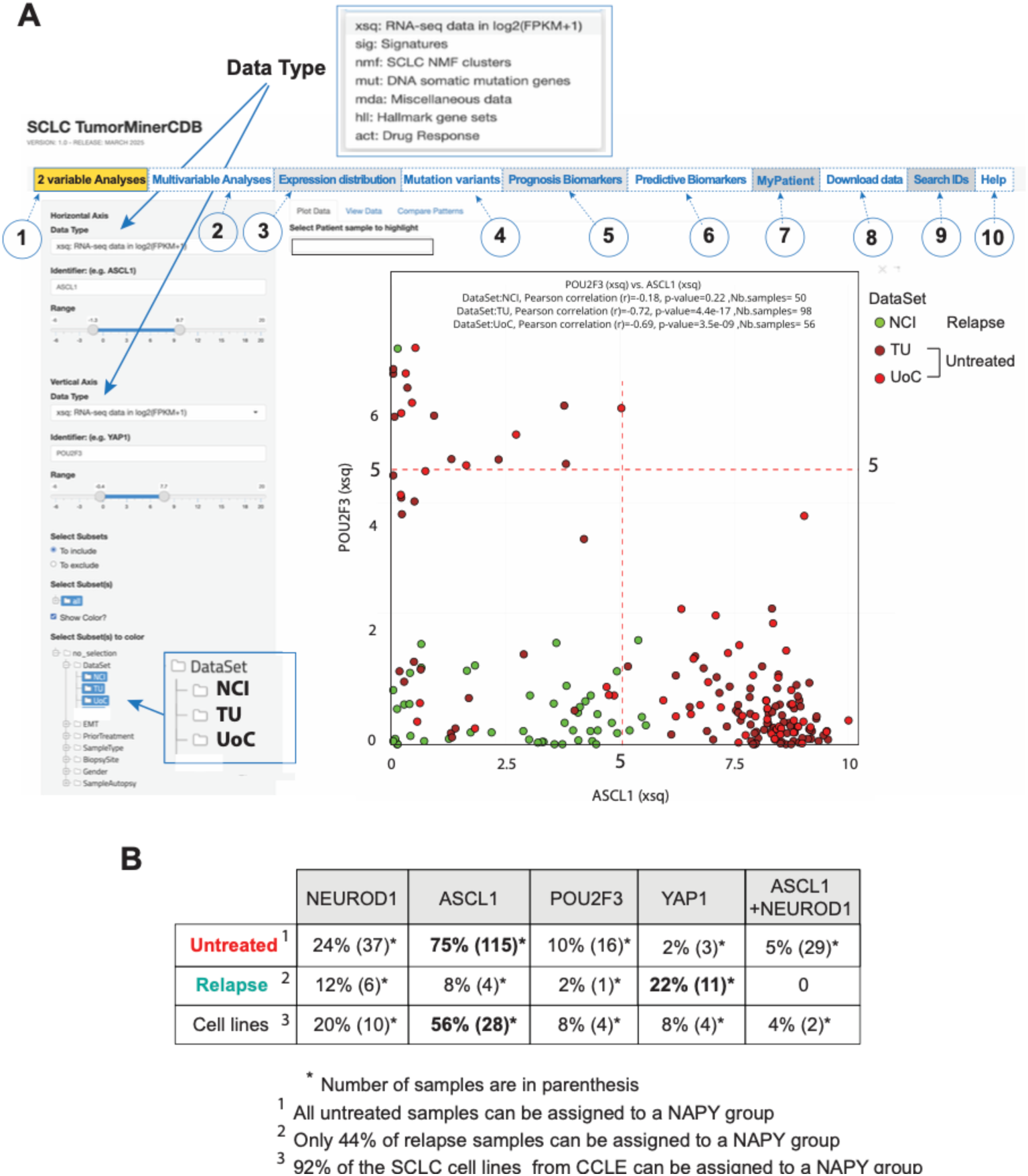
Introduction to SCLC TumorMiner (https://discover.nci.nih.gov/SclcTumorMinerCDB/). **A**. Snapshot of the introductory “2 variable Analyses” display (<u>1</u>). Both the horizontal and vertical axes offer a choice of “Data Type” shown in the upper inset. Users must type their choice in the “Identifier” box (here ASCL1 for the horizontal axis and POU2F3 for the vertical axis). Other options include choosing the “Range” for the axes and the “Select Subsets” to “include” or “exclude”. The + sign and folder indicate that further selections can be made such as choosing the “Dataset” to display. To display more than one dataset, users need to press the Command key while clicking the individual datasets (lower inset). Other features of SCLC TumorMiner include: (<u>2</u>) “Multivariable Analyses” with examples shown in Figure 2; (<u>3</u>) “Expression Distribution” across the 4 Datasets with example shown in Figure S1; (<u>4</u>) “Mutation variants”; (<u>5</u>) “Prognosis Biomarkers” with example shown in Figures 5 and S4; (<u>6</u>) “Predictive Biomarkers” with example shown in Figure 6; (<u>7</u>) “My Patient” to display the genomic characteristics of a selected patient with example shown in Figures 7 and S6. Additional features include the ability to “Download Data” for further analyses (<u>8</u>) as shown in Figures 7, S1, S4 and S5, “Search IDs” (<u>9</u>), as well as a detailed “Help” section (<u>10</u>). **B**. NAPY distribution of samples from untreated and relapse patients and from the SCLC cell lines (https://discover.nci.nih.gov/SclcCellMinerCDB) ^4^. Samples were assigned to each NAPY group based on gene expression log2(FPKM+1) > 5.0 (see horizontal and vertical dotted lines in panel A and justification for the > 5.0 threshold in Figure S1).

Immune checkpoint (PD-L1) inhibitors added to chemotherapy are the standard of care for first-line treatment of ES-SCLC ^6,7^. However, they only provide a modest overall survival benefit of 2-3 months ^7,8^. A recent therapeutic breakthrough for SCLC is Tarlatamab^®^, a bispecific T-cell engager (BiTE^®^) targeting the Notch ligand cell surface glycoprotein DLL3 (Delta-like ligand 3), a transcriptional target of ASCL1 ^4^. However, responses are heterogeneous, and Tarlatamab^®^ is currently administered without mandatory biomarker-based patient selection.

In parallel, antibody–drug conjugates (ADCs) and CAR-T therapies targeting surface antigens such as DLL3, TROP2 (anti-trophoblast antigen 2 encoded by *TACSTD2*), SEZ6 (seizure 6-like), carcinoembryonic antigen–related cell adhesion molecules (CEACAM5 and CEACAM6), and B7-H3 (member of the B7-cell family homolog 3 encoded by *CD276*) are actively being developed. For ADCs in particular, therapeutic efficacy depends not only on target expression but also on payload sensitivity, including replication stress–associated vulnerabilities such as expression of Schlafen 11 (SLFN11) and Ki67 (encoded by *MKI67*).

With the advancement of high-throughput genomic sequencing methods, bulk RNA sequencing (RNA-seq) has emerged as a well-established, sensitive, and clinically usable method for transcriptome analyses. However, even when only applied to protein-coding genes, it generates around 20,000 gene expression data per sample, which are difficult to analyze instantaneously in a clinical setting. To address this challenge, we developed SCLC TumorMiner, integrating 204 SCLC patient samples. Fifty samples are collected from patients at relapse in our NCI clinic, and 154 are from the published datasets of untreated patients from the University of Cologne (UoC) ^2^ and Tongji University (TU) ^9^ (Table 1, Table S1). The architecture and interface of TumorMiner is based on our SCLC CellMinerCDB website comprising the genomics and drug responses of 118 SCLC cell lines (https://discover.nci.nih.gov/SclcCellMinerCDB) ^4^. Thus, SCLC TumorMiner and SCLC-CellMinerCDB can be readily used to compare tumor samples with cell lines.

**Table 1.** SCLC samples and Datasets

|  |  | Untreated |  | Relapse | Total |
| --- | --- | --- | --- | --- | --- |
|  |  | UoC | TU | NCI |  |
| **Number of Samples** |  | 56 | 98 | 50 | 204 |
| **Genomics** | RNA Seq | 56 | 98 | 50 | 204 |
|  | WES | 0 | 98 | 16 | 114 |
| **Prior Treatment** | Yes | 0 | 0 | 50 | 50 |
|  | No | 52 | 98 | 0 | 150 |
|  | Unknown | 4 | 0 | 0 | 4 |
| **Biopsy Site** | Lung | 55 | 98 | 7 | 160 |
|  | Liver | 0 | 0 | 21 | 21 |
|  | Lymph Node | 0 | 0 | 10 | 10 |
|  | Pleura | 1 | 0 | 1 | 2 |
|  | Adrenal Gland | 0 | 0 | 4 | 4 |
|  | Mediastinal mass | 0 | 0 | 3 | 3 |
|  | Others | 0 | 0 | 4 | 4 |
| **Tumor Stage** | Stage 1 | 25 | 30 | 0 | 55 |
|  | Stage 2 | 7 | 26 | 0 | 33 |
|  | Stage 3 | 17 | 41 | 2 | 60 |
|  | Stage 4 | 7 | 1 | 48 | 56 |

Here we provide a tutorial and demonstrate how SCLC TumorMiner allows instantaneous: 1/ Transcriptome and mutations analyses of patient samples; 2/ Molecular pathway analyses; 3/ Patient sample classification based on NAPY and/or NMF (non-negative matrix factorization); and 4/ Identification of prognosis and predictive biomarkers for targeted therapies.

## RESULTS

### Introduction to SCLC TumorMiner and brief tutorial

SCLC TumorMiner uses the same architecture as our six-cancer cell line CellMiner websites (https://discover.nci.nih.gov/). Thus, users familiar with SCLC-CellMinerCDB (https://discover.nci.nih.gov/SclcCellMinerCDB/) will have no difficulty navigating back and forth between the tumor samples and cell lines databases ^4^.

Figure 1 (https://discover.nci.nih.gov/SclcTumorMinerCDB/) introduces the website as a snapshot of the default page display. The upper part shows the tabs for each of the tools and applications. They are numbered 1-10 for convenience on the figure. The “2 variable Analyses” tab (numbered “1” in Figure 1A) allows the display of different data types or parameters on each of the axes: “xsq” (RNA-seq), “sig” (Signatures), “mut” (gene mutations), “mda” (Miscellaneous data), “hll” (Hallmark gene sets) and “act” (Drug Response). For instance, plotting ASCL1 expression vs. NE (Neuro Endocrine) signature (Figure S1A) shows their expected high correlation within each of the 3 datasets (NCI, UoC, TU).

2-Variable Analysis data can be downloaded as Tab-delimited files for further exploration. Figure 1B compares the expression of *NEUROD1*, *ASCL1*, *POU2F3* and *YAP1* across datasets. It demonstrates the reduced *ASCL1* and *POU2F3* expression associated with increased *YAP1* expression in the relapse samples (Figure 1A). As a result, the assignment of samples to each of the NAPY groups can only be applied to 44% of the relapse samples while, it discriminates all the samples from untreated patients ^3^ and 92% of the cell lines from SCLC-CellMinerCDB ^4^ (Figure 1B).

The “Multivariable Analyses” tab (labeled “2” in Figure 1A) shows as default display the heatmap of *ASCL1* expression (xsq: RNA-seq) vs. the expression of key genes driving SCLC: *INSM1* & *CHGA* encoding immunohistochemistry NE markers, *MKI67* (encoding Ki67 used as proliferation marker), and the SCLC pioneer transcription factors *NEUROD1*, *POU2F3* & *YAP1*, and the oncogene *MYC* (Figure 2A). Each row displays the selected genes, and the columns represent each of the 204 individual patient samples. Users can explore different inputs and outputs by entering gene names in the “Response Identifier” or “Predictor Identifiers” windows (left panels of Figure 2A). Examples of “Multivariate Analyses” for the NAPY genes are shown in Figure 2A-E and will be discussed in the next section.

**Figure 2.**
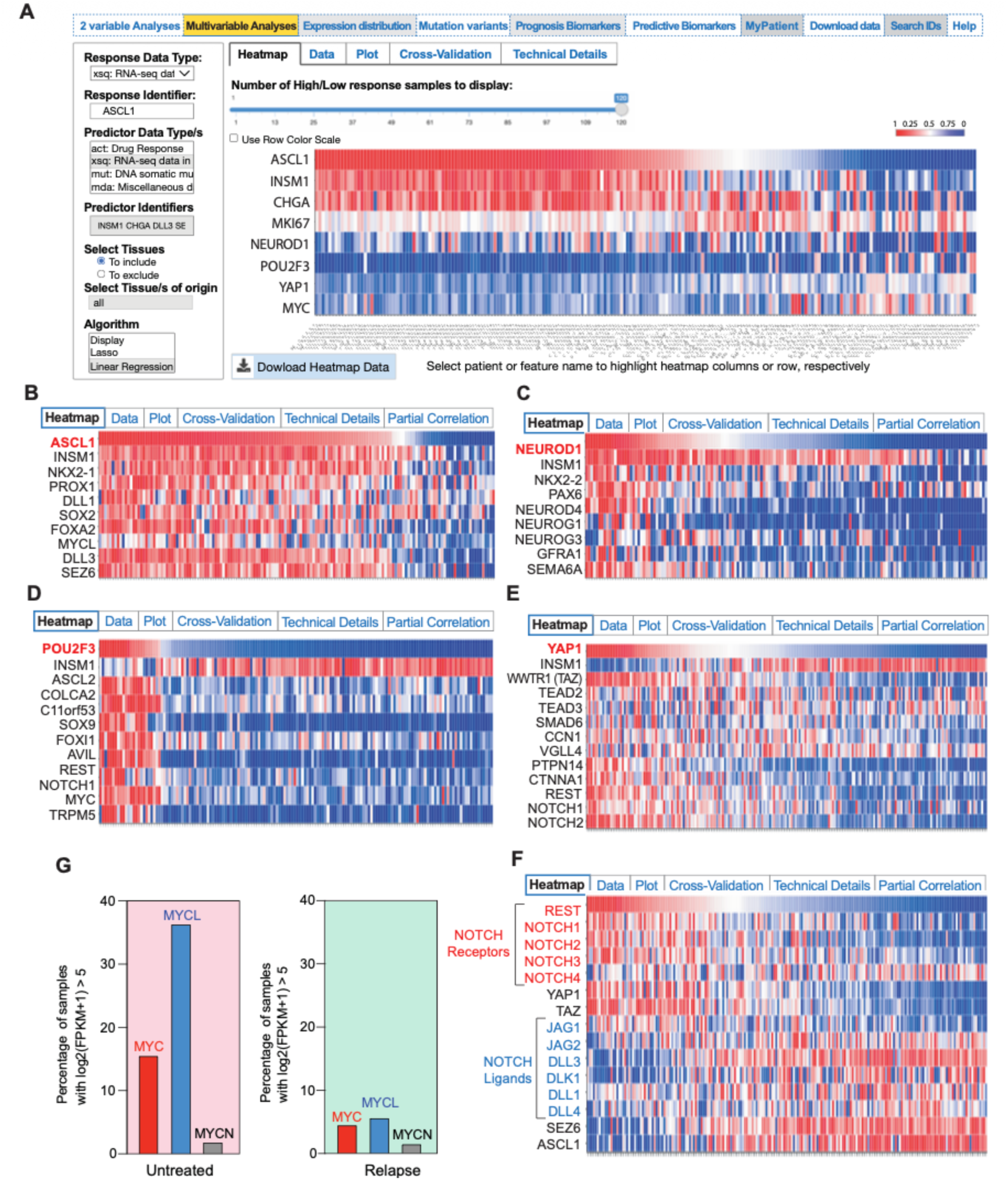
Multivariable Analyses showing co-regulation of the neuroendocrine (INSM1, CHGA), NAPY (NEUROD1, ASCL1, POU2F3 and YAP1), NOTCH (NOTCH1, NOTCH2, REST) and MYC (MYC, MYCL and MYCN) pathways. **A**. Default display for the “Multivariable Analyses” tab. Color scale goes from red to blue to reflect expression levels for each gene (rows) and patients (columns). Users can choose different data types (upper inset of panel A) and then type the “Response Identifier” to display as the first row on the heatmap. “Predictor Identifier (s)” need to be typed with a space between each. “Tissues of origin” can be selected. The default algorithm (bottom left) is “Display”. Users can also perform Linear and Lasso analyses like in CellMinerCDB (https://discover.nci.nih.gov/cellminercdb/). **B-E**. Multivariable analyses for the NAPY genes used as “Response Identifiers”. I*NSM1* is included in each analysis as a NE biomarker. Panels B-D only include the samples from the databases of untreated patients. **F**. Multivariable analysis for the Notch pathway. **G**. Differential expression of the 3 MYC genes in samples from untreated and relapse patients. Linear regression analyses for each of “Response Identifiers” (in red, 1^st^ rows in panels B-F) show that the “Predictor Identifiers” are highly significant (see Figure S2C).

Under the “Multivariate Analyses” tab, in addition to the default visualization set up by default as “Display”, users can choose the “Linear Regression” tab (found at the bottom left corner of the website under “Algorithm”; Figure 2A). This feature computes the correlation between each of the “Predictive Identifiers” and the “Response Identifier”. Figure S2A-B shows the results for *ASCL1* for the genes entered in Figure 2B. The “Plot” and “Technical Details” options illustrate the highly significant value of the combined expression of *NKX2.1*, *FOXA2*, *DLL3* and *SEZ6* as predictors of *ASCL1* expression in the combined datasets (UoC and TU) of the untreated patients (Figure S2A-B). Users can also choose “Lasso” (Least Absolute Shrinkage and Selection Operator), a machine-learning and statistical method to identify genomic biomarkers predicting the expression of “Response Identifiers”. For *YAP1* expression, the top gene identified in the relapse samples is the Neuron Restrictive Silencing Factor *REST*, a key effector of the Notch pathway (Figure S2D) ^4^.

The “Expression Distribution” tab (labeled “3” in Figure 1A) displays the expression of any chosen gene across the 3 datasets (the two untreated TU & UoC datasets and the relapse patients from the NCI dataset). *SLFN11* expression is displayed as default on the website (Figure S3A). Its reduced expression in the relapse samples is consistent with the resistance of relapsing patients to DNA-targeted chemotherapies. Users can also display and download data for any gene from each of the 3 datasets by typing the gene HUGO name in the “Gene symbol” box. Such analyses for the housekeeping genes *GADPH*, *ACTB* (β-actin), or *B2M* (β2-macroglobulin) show high expression and lack of significant expression differences between untreated and relapse samples, suggesting limited batch effect for highly expressed genes (>5 in at least one dataset).

The “Mutation Variants” tab (labeled “4” in Figure 1A) displays up to 5 genes at a time. Colors reflect the different types of mutations, including: Missense_Mutation, Nonsense_Mutation, Frame_shift_Ins, Frame_Shit_Del, In_Frame_Del, Multi_Hit. Variants can be viewed and downloaded with the “View variants” tab (Figure S3B). As expected, *TP53* is the most frequently mutated gene (81% of the 114 samples with whole-exome sequencing data). The second most frequently mutated gene is *TTN,* encoding Titin. This is not surprising as Titin is the largest human protein and SCLC has a high mutation burden. RB1 shows an overall mutation frequency of 55% (Figure S3B). By computing the copy number (Ploidy) values for RB1 for the TU and NCI SCLC (114) samples and using an *ad hoc* rule (samples with < 0.05 ploidy for full depletion and between 0.05 and 1.85 for partial depletion), percentages for full and partial depletions are 62.8% and 17.7%, respectively (Table S2). Inactivation of SLFN11 was observed in only 4% of samples (Figure S3B), consistent with epigenetic inactivation as the main mechanism for suppressing SLFN11 expression and conferring chemoresistance ^10^.

The “Prognosis Biomarkers”, “Predictive Biomarkers” and “MyPatient” tabs (numbered 5, 6 & 7 in Figure 1A) will be discussed in the remaining sections of the manuscript with examples displayed in Figures 5 & 6.

**Figure 3.**
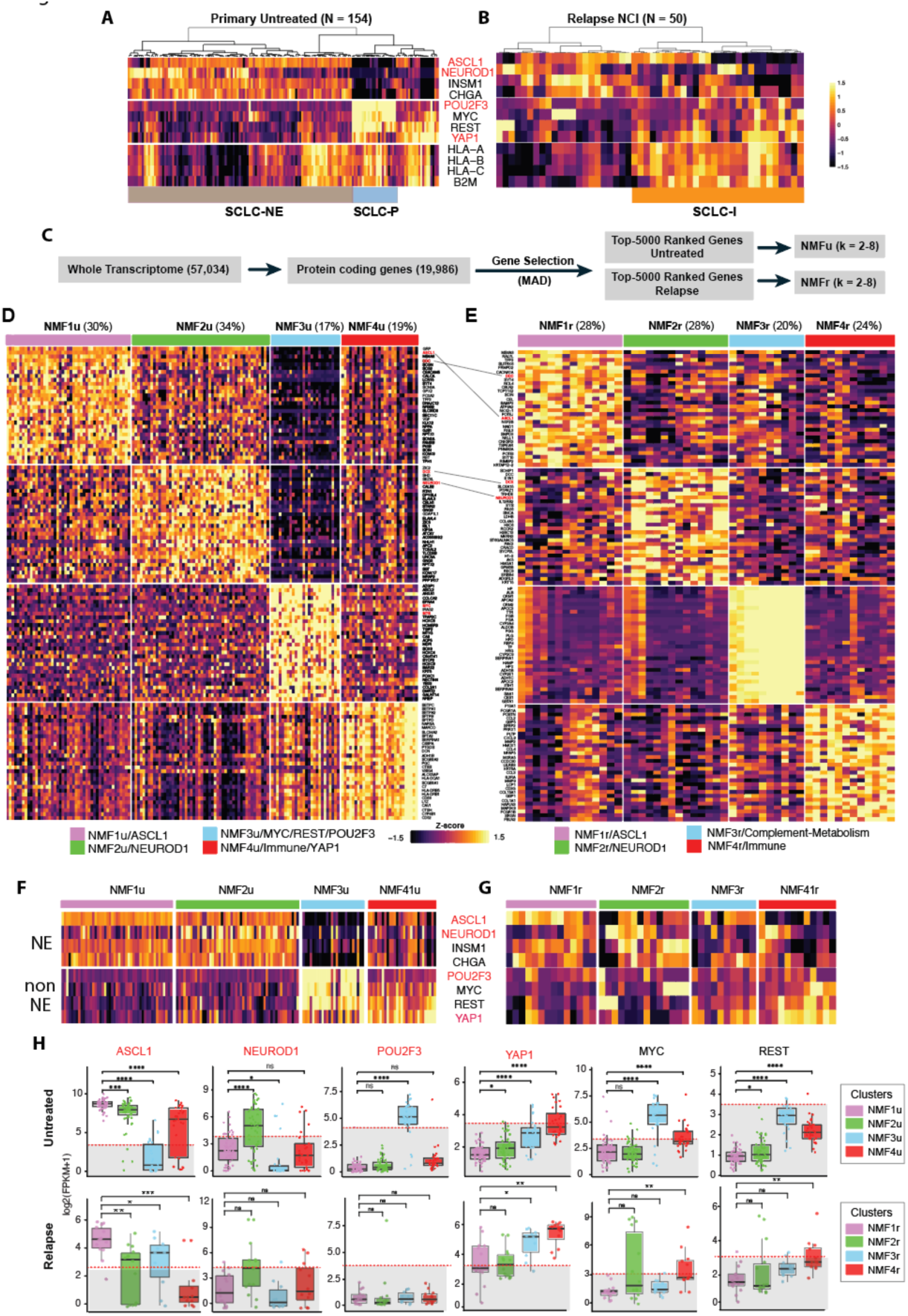
Classification of the untreated (primary) and relapse 204 SCLC samples comparing untreated (UoC and TU datasets) and relapse samples (NCI). **A-B**. Hierarchical clustering of the untreated and relapse samples based on the expression of the four genes of the NAPY classification (*NEUROD1*, *ASCL1*, *POU2F3* and *YAP1* in red) and of genes reflecting commonly used biomarkers for clinical neuroendocrine (NE) classification (*INSM1* and *CHGA*) as well as *MYC*, *REST* and 4 exemplary innate immune genes (*HLA-A*, *HLA-B*, *HLA-C* and *B2M)*. **C**. Flow chart summarizing the NMF approach for selecting 5,000 protein-coding transcripts based on MAD (Median Absolute Deviation) for the untreated and relapse samples. **D-E**. Display of the NMF clusters for the untreated and relapse samples (D and E, respectively). The percentages of samples within each NMF are indicated in parenthesis. NMFs for untreated and relapse are abbreviated NMFu and NMFr, respectively. The rows show the top 30 genes for each NMF with log fold-change > 1 and p-value < 0.05. Genes that are common across untreated and relapse samples for the NMF clusters are indicated by the oblique lines. **F-G**. NMF distribution of the canonical NAPY genes (red) and reference NE genes as well as *MYC* and *REST.* **H**. Box plots comparing the differential expression of the NAPY genes, *MYC* and *REST* across the 4 untreated (top) and relapse NMFs (bottom). Statistical comparisons were performed by Student t-test. **** indicates p < 0.001; ** p < 0.01; * p < 0.05. Shaded areas represent low expression (log2(FPKM+1) <3).

**Figure 4.**
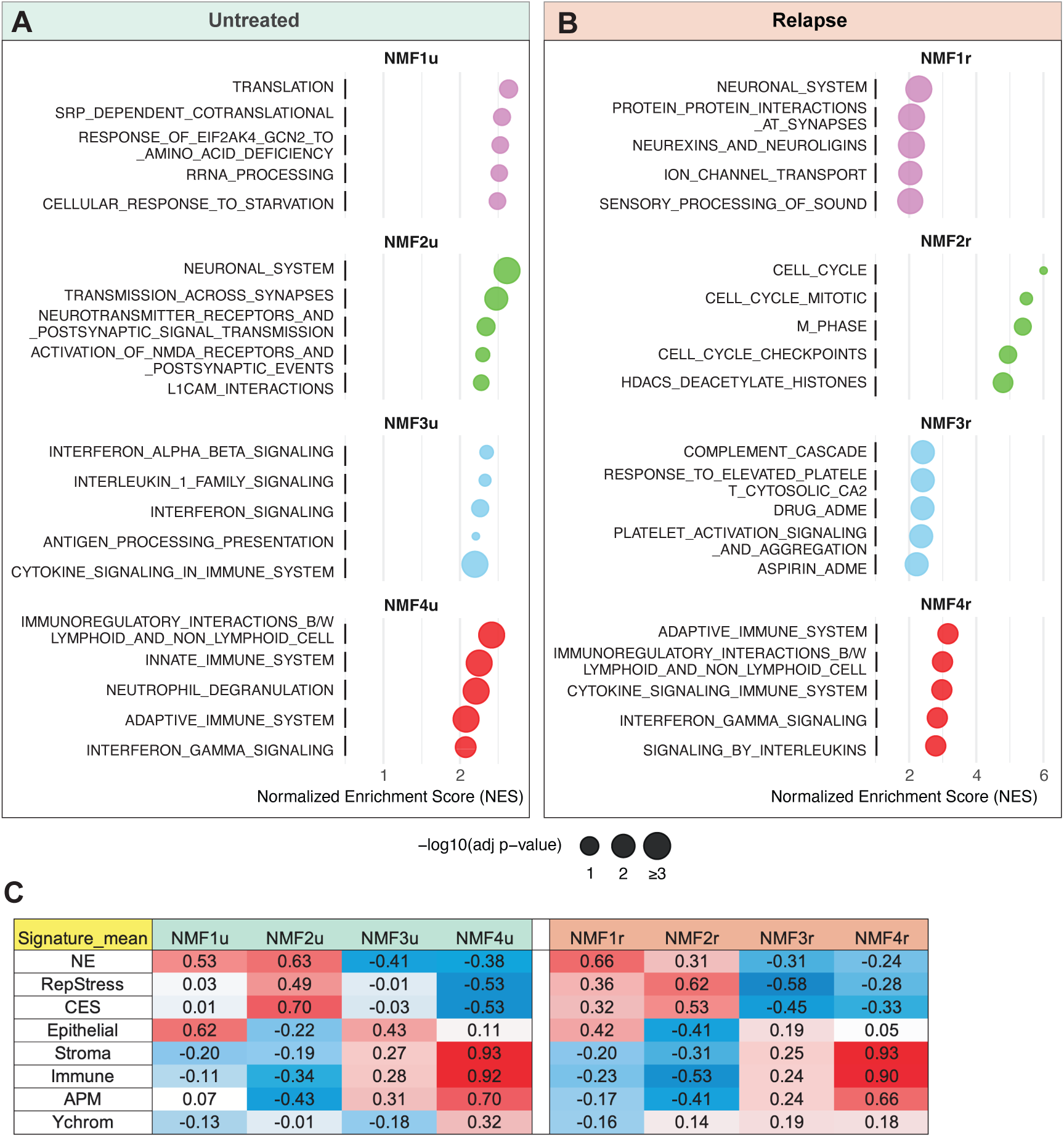
Genomic signatures associated with each NMF from the untreated and relapse patient biopsy samples. **A-B**. Reactome pathway analyses for untreated and relapse samples. **B**. Correlation between each of the NMF’s and the eight genomic signatures of SCLC TumorMiner for the samples from untreated (left) and relapse (right) patients. Correlations were obtained using the “2 variable Analyses” of the SCLC TumorMiner website (“Plot Data” or “View Data” tabs for each of the NMF on the horizontal axis). For the untreated samples the corresponding NMF’s were computed after limiting the analysis using “Select Subsets” for the untreated Datasets: TU + UoC. Conversely for the relapse samples, analyses were done using the NCI samples:

**Figure 5.**
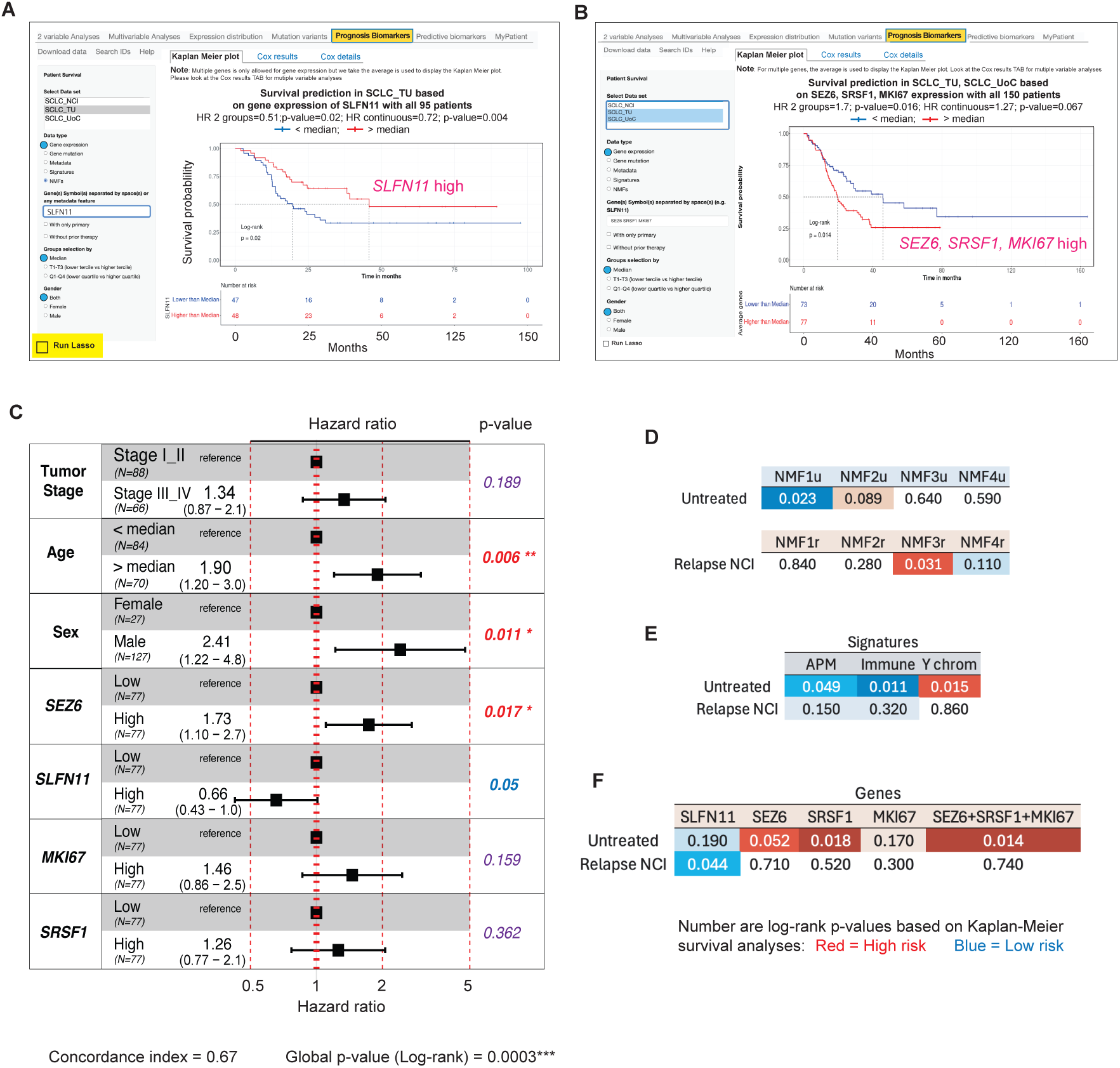
Exploration of prognosis biomarkers. Panels A and F are screen snapshot of the SCLC TumorMiner website. **A**. High *SLFN11* expression is a good prognosis biomarker for the untreated TU Dataset. **B**. Combined expression of *SEZ6*, *SRSF1* and *MKI67* are significant risk factors for the combined untreated datasets (TU & UoC). **C**. Multivariate survival analysis including *SEZ6, SLFN11*, *SRSF1*, *MKI67* expression, age, tumor stage and sex for the untreated patients (TU and UoC datasets). **D**. Tabulated results for the prognosis value of each NMF cluster. **E**. Prognostic value of 3 of the 8 signatures of SCLC TumorMiner. **F**. Prognostic value of *SLFN11*, *SEZ6, MKI67* and combined expression of *SEZ6*, *SRSF1* and *MKI67* as prognosis biomarkers for the untreated and relapse datasets. Numbers (panels D-F) are log-rank p-values based on Kaplan-Meier survival analysis and colors highlight significant value (p < 0.05) with low risk parameter in blue and high-risk in red.

### Genomic networks and pathway explorations

We previously reported ^4^ in the SCLC cell lines, the expression of genes of a given pathway are cross-correlated [see Fig. 5 in ^4^]. The main pathways for SCLC comprise the neuroendocrine (NE) pathways driven prominently by *ASCL1* and less frequently by *NEUROD1*, and the non-NE pathways driven by *POU2F3* and *YAP1*, the *MYC* genes, and the Notch pathway ^3,4^. As in the SCLC cell lines (https://discover.nci.nih.gov/SclcCellMinerCDB/) ^4^, the group of NE-SCLC prominently expresses *INSM1* and *CHGA,* encoding routinely used immunochemistry biomarkers (Figure 2A). As expected, *ASCL1* expression is highly correlated with *INSM1* and *CHGA*. Figure 2A also shows that multiple samples express both *ASCL1* and *NEUROD1*, and a significant overlap between *YAP1*, *POU2F3* and *MYC* expression.

Analysis of the *ASCL1* coexpression network in the samples from untreated patients (Figure 2B) confirms ^4^ that most samples expressing *ASCL1* coexpress *INSM1* (highly significant correlations in the “2 variable analysis”). As expected, the transcriptional coactivators of *ASCL1*, *NKX2-1* and *PROX1* ^4,11^ show coexpression (confirmed in the “2 variable analysis”). *DLL3*, the downstream transcriptional target of ASCL1, is highly expressed in approximately two-third of the samples with high correlation with *ASCL1* expression (confirmed in the “2 Variable Analyses”). *SEZ6*, another ASCL1 target gene and an emerging cell surface target for Antibody Drug Conjugates is also highly correlated with both *ASCL1* and *DLL3* expression (Figure 2B). Of note, *SEZ6* expression is highly correlated with *SEZ6L* (SEZ6 homolog like) despite different chromosome locations (17q11.2 and 22q12.1, respectively), suggesting that both genes function in the ASCL1 pathways.

Transcriptome analysis of the NEUROD1 genomic network (Figure 2C) shows that *NEUROD1* expression only accounts for a fraction of the NE samples. Indeed, many samples show high *INSM1* expression without high *NEUROD1* (Figure 2C, compare the 1^st^ and 2^nd^ rows). *NKX2-2* [NK2 transcription factor related, locus 2 (Drosophila)], which is involved in the morphogenesis of the central nervous system (CNS) and a known regulator of NEUROD1 in pancreatic islet cells together with the Neurogenin genes (*NEUROG1* and *NEUROG3*) ^12^ are co-expressed with *NEUROD1* (Figure 2C). The coexpression of *PAX6* (Paired Box Homeotic Gene-6) encoding a transcription factor for the development of the eyes, nose, pancreas and CNS, is logical as PAX6 controls *NEUROD1* expression ^13^. Using the “2 variable Analysis” and the “Compare Patterns” tools of SCLC TumorMiner (see Figure 1A), we discovered coexpression of two genes encoding cell surface proteins and potential targets for immune therapies, *GFRA1* (Glial Cell Line-Derived Neurotrophic Factor Receptor Alpha) and *SEMA6A* (encoding one of the semaphorin) ^14^ (Figure 2C). Like the samples from untreated patients, the SCLC cell lines also express significant levels of *SEMA6A* with high correlation with *NEUROD1* (p = 0.57 with p-value = 2.7e-11) ^4^. Whether NEUROD1 directly drives *GFRA1* and *SEMA6A* expression remains to be determined.

The POU2F3 network shows the expected inverse expression between *POU2F3* and *INSM1* (Figure 2D, compare the 1^st^ and 2^nd^ rows) as the POU2F3 cells are derived from the Tuft cells of the bronchial epithelium and are non-NE ^15–17^. Figure 2D shows that the tumor samples expressing *POU2F3* coexpress 4 transcription factors known to drive *POU2F3* expression ^15,18^: *ASCL2, COLCA2* (also known as *OCAT-2* or *POU2AF3*), *C11orf53* (also known as *OCAT-1* or *POU2AF2*) and *SOX9*. *POU2F3* is also highly correlated with the Tuft-ionocyte progenitor specific genes *FOXI1* and *AVIL* ^16,17^, as well as with *TRPM5* (Transient Receptor Potential Cation Channel Subfamily M Member 5), which encodes a cell surface ion channel protein highly expressed in bronchial Tuft cells and involved in immune defense ^19^. This suggests that TRPM5 could be considered as cell surface target for SCLC-P tumors. Figure 2D also shows that the POU2F3 network/pathway is coordinated with the Notch pathways, as *POU2F3* expression is highly correlated with *REST* and its transcriptional activator *NOTCH1* (validated in the “2 variable analysis” for the untreated samples). *MYC* expression is also highly correlated with *POU2F3*, consistent with independent reports ^9,17,20^. These findings suggest that the expression of the 11 genes displayed in Figure 2D could be used as predictors for POU2F3 expression in tumor samples (Figure S2C) and for classifying a given tumor sample as SCLC-P.

Analysis of the YAP1 network across our 3 datasets shows that samples with high *YAP1* expression tend to lack *INSM1* expression (Figure 2E). By contrast, TAZ (encoded by *WWTR1*), the transcriptional co-activator partner of YAP1 is coexpressed with *YAP1*, consistent with the SCLC cell lines (Figure 5 in ^4^). Similarly, *TEAD2* and *TEAD3*, the downstream transcription factors of YAP/TAZ are coexpressed with *YAP1*, as well as YAP1’s transcriptional targets *SMAD6*, *VGLL4* and *PTPN141* ^4^. Figure 2 also shows that the cell adhesion genes *CCN1*, *CTNNA1* and *CTNNA2* are coexpressed with *YAP1*. Consistent with the SCLC cell line data (see Figure 5C-D in ^4^), Notch pathway activation (*REST*, *NOTCH1* and *NOTCH2*) is associated with the SCLC-Y samples. Together the expression of the 12 genes displayed in Figure 2E significantly predicts *YAP1* expression both in the SCLC tumors and cell lines (Figure S2C) and therefore could be used to classify a given tumor sample as SCLC-Y (Figure S2D).

Using *REST* expression as a reporter for the Notch pathway (Figure 2F), we found coexpression of *REST* with the 4 Notch receptors, *NOTCH1-4* and with YAP/TAZ, consistent activation of the Notch pathway by YAP1-mediated transcription ^21^. By contrast, expression of the Notch ligands (*JAG1/2*, *DLL3*, *DLL1*, *DLL4* and *DLK1*) is significantly suppressed in the Notch receptor-positive samples and coexpressed with *DLL3* and *ASCL1* (Figure 2F). These findings support the conclusion that the YAP/Notch/REST network controls the NE fate of SCLC cells ^21^ and vice-versa as DLL3 suppresses Notch signaling in SCLC ^22^.

Figure 2G shows MYC pathway activation in 52% of the untreated patient samples where *MYCL* dominates (36.4% of the samples). *MYCL* expression also appears linked with *ASCL1* expression (SCLC-A) (Figure 2B). By contrast, *MYC* is only highly expressed in a relatively small fraction of the untreated patient samples (15.6%) and primarily in the SCLC-P samples (Figures 2D) ^23^. Consistent with the fact that *POU2F3* disappears in the relapse samples and that *ASCL1* expression is also attenuated in those samples, the expression of both *MYCL* and *MYC* appears markedly reduced in the samples of patients at relapse (Figure 2G).

The conservation of each of the 5 networks described above between tumor samples and cell lines (Figure S2C) implies their predictive value as genomic signatures for tumors driven by ASCL1, NEUROD1, POU2F3, YAP1 and Notch.

### NAPY classification of SCLC tumors

To identify highly expressed genes, we first analyzed the gene expression distribution for all genes across the datasets (Figure S1B). The 90-percentile expression value cutoff selected log2(FPKM+1) values > 4 for both the untreated and relapse samples. To further increase stringency, we set up a threshold value log2(FPKM+1) > 5 for high gene expression (red lines in Figures 1A and S1A-C, and shaded areas in Figure S1B-C). We also compared the expression of canonical housekeeping genes in the untreated and relapse datasets and found their expression well-above the threshold of 5 and relatively comparable between the untreated and relapse datasets, suggesting limited batch effect (Figure S1C).

Using RNA-seq expression levels >5 [log2(FPKM+1)], we could unambiguously assign all the samples from the untreated patients to the NAPY classification (Figures 1B). However, less than half (44%) of the samples from relapse NCI patients could be classified with 22% as high *YAP1* (Figure 1B). This is notably higher than for the untreated patients (22% vs. 2% *YAP1* high; Figure 1B). Notably, based on the same cut-off [log2(FPKM+1)>5], the SCLC cell lines could be classified like the untreated samples, which is consistent with the fact that most SCLC cell lines have been derived from patients before treatment ^24,25^.

To classify the samples from relapse patients, we first attempted to extend the gene list for cluster analyses by adding to the 4 NAPY genes (highlighted in red in Figure 3A-B), the two NE genes (*INSM1* and *CHGA*), *MYC*, *REST,* and 4 innate immune genes of the major histocompatibility complex (MHC): *HLA-A*, *HLA-B*, *HLA-C* and *B2M* as reporters of an inflamed signature (SCLC-I) ^5^. This 12-gene algorithm classified the samples of untreated patients in 2 groups (Figure 3A; see annotations under the heatmap): SCLC-NE (≈ 70% of the samples) and SCLC-P with high *POU2F3*, *MYC*, *REST* and to a lesser extent *YAP1 e*xpression. The inflamed group with high MHC expression (*HLA-A/B/C* and *B2M*) was found diffuse (≈ 50% of the patient samples) with tendency to be excluded from the SCLC-NE group. For the samples of relapsed patients, the 12 gene signature only clustered the samples based on the MHC genes (≈ 50% of the samples; SCLC-I) (Figure 3B). To overcome the difficulty of classifying the samples from patients at relapse using the NAPY classification, we performed NMF (Non-Negative Matrix Factorization) analyses ^5^.

### NMF classification of SCLC tumors

To perform these NMF analyses, we used the top 5000 most variably expressed protein-coding genes separately in the untreated (UoC and TU) and relapse (NCI) datasets (Figure 3C, Tables S3 and S4). Among those genes, 64.4% are common for the untreated and relapse samples (Figure S4A). Cophenetic and Silhouette scores showed that 4 clusters gave a satisfactory division of the samples (Figure S4B-C).

To develop open-access NMF signatures, we first took the genes with a 2-fold selectivity compared to the genes in the other 3 NMF groups. This led to a list of genes in each NMF group. For the relapse samples, we selected 181 genes for NMF1r, 58 for NMF2r, 342 for NMF3r and 215 for NMF4r (Table S5). For the untreated samples, we obtained 54 genes for NMF1u, 137 for NMF2u, 85 for NMF3u and 386 for NMF4u (Table S6). At the end, we obtained 662 and 795 genes for the untreated and relapse cohorts, respectively (see Method section for details). The top-30 genes for each of the NMFu and NMFr clusters are displayed in Figure 3D and Figure 3E, respectively. Notably, only a few genes are common in the NMF clusters of untreated and relapse patient samples (see diagonal lines between panels D and E in Figure 3 and see Tables S5 and S6).

Pathway analysis shows that NMF1u consists of genes involved in translation and RNA metabolism (Figure 4A) in addition to *ASCL1* (Figures 3D, F, H)). The NE enrichment of NMF1u is easily visible through the mapping *ASCL1*, *NEUROD1*, *INSM1* and *CHGA* in the NMFu clusters (Figure 3F). NMF2u also shows strong NE enrichment (Figures 3F and 4A) and includes *NEUROD1* (Figure 3D). NMF3u includes *POU2F3* (Figures 3F and 3H) and immune and interleukin pathways (Figure 4A) as well as the MYC pathway (Figures 3D; *MYC* and *MYB* annotated in red). NMF4u partially extends to NMF3u and shows positive enrichment for immune, cytokine and interferon pathways (Figures 3D and 4A). It is enriched for *YAP1* (Figures 3F & H).

Pathway analyses for the 4 relapse NMF clusters shows that NMF1r includes genes and pathways related to NE differentiation (Figures 4B and 3E) as well as *ASCL1* (Figures 3G-H). NMF2r comprises NEUROD1 (Figure 3E and 3H) and is enriched for cell cycle genes (Figure 4B). NMF3r includes genes involved in metabolism and complement (Figure 4B). It also includes *YAP1* (Figure 3G-H). NMF4r consists specifically of immune genes (Figures 3E and 4B) and some *YAP1*-expressing samples (Figures 3G and 3H). The complete pathway enrichment results for untreated and relapse SCLC samples are provided in Table S7 and Table S8, respectively.

### The NMF clusters reflect the SCLC TumorMiner molecular signatures

Next, we explored the correlations between the NMF and eight transcriptional signatures provided in SCLC TumorMiner: NE (Neuroendocrine), RepStress (Replication stress), CES (Chromosome Centromere and Kinetochore score), Epithelial, Stroma, Immune, APM (Antigen Presenting Machinery) and Ychrom (Y-chromosome), which were computed as the mean expression of the genes included in each signature (see Methods section).

Figure 4C tabulates the Pearson coefficient correlations between each NMF and the 8 signatures with the colors reflecting the direction and intensity of the correlations (red for positive and blue for negative). Consistent with the top gene signatures and the Reactome pathway analyses (see Figure 4A-B), the NMF1u, NMF2u, NMF1r and NMF2r signatures are highly correlated with the NE signature as well as with the RepStress and CES signatures. They are also significantly devoid of the Immune, APM and Stroma signatures (Figure 4C). Conversely, NMF3u, NMF3r, NMF4u and NMF4r are enriched for the Immune, APM, Stroma and epithelial signatures and lack the NE, RepStress and CES signatures. Together these results demonstrate the functional significance of the NMF classifications for SCLC.

### External validation of the NMFs

To evaluate and validate our NMF-based gene signatures using independent datasets, while avoiding batch effect correction with the training data, we computed NMF scores for each sample based on the signature genes. We then clustered samples using the top 30 most differentially enriched genes and assessed whether the resulting patterns were consistent with those observed in the training datasets. We first performed validation using RNA-seq data from two external untreated SCLC cohorts, GSE60052 ^26^ (n = 79) and IMpower133 ^8^ (n = 271). As both cohorts consist of treatment-naïve samples, we applied the NMFu untreated signatures for clustering and validation (Table S6). As shown in Supplementary Figure S5A-B, the top 30 most differentially enriched genes were consistent with those identified in our untreated training SCLC cohort. Neuroendocrine (NE) markers, including ASCL1, NEUROD1, INSM1 and CHGA, as well as non-NE markers such as POU2F3, YAP1, MYC and REST, exhibited distinct and consistent enrichment patterns across clusters in both external cohorts. These findings support the biological relevance and reproducibility by identifying the same 4 NMF signatures for untreated samples in these two other external datasets.

Next, we sought to validate the NMF relapse signature using the Wagner dataset ^27^, which includes 22 samples from 18 relapse SCLC patients. As this dataset contains two technical batches (RNA-seq protocols: polyA and total RNA; both derived from frozen samples), we applied batch correction as described in the original study. As shown in Figure S6A, the top 30 most differentially enriched genes were consistent with those observed in three of our relapse SCLC clusters: NMF1r, NMF2r and NMF4r. We did not identify a cluster corresponding to NMF3r of the NCI training dataset, which is associated with complement and metabolic pathways. To further investigate the consistency of the 4 NMF relapsed clusters, we applied the NMF method on the Wagner dataset using the same set of input genes (∼5000 genes) (as with the NCI dataset) and computed the correlation between the metagenes from the NCI basis matrix to the equivalent matrix generated for the Wagner dataset. Figure S6C shows the correlation of Wagner metagene groups with the NCI metagene groups. Wagner W3 group only had low correlation to the NCI groups (r=0.29), whereas other subgroups had correlations greater than 0.6 (for W1, W2 and W4). This discrepancy may be due to differences in sample composition, including the absence of liver metastasis samples and variation in biopsy sites between the NCI and Wagner datasets (see Figure S6B).

### Prognosis biomarkers and risk factors

SCLC TumorMiner is designed to assess prognosis biomarkers associated with gene expression, mutations, clinical metadata and NMF signatures. The default display on the website is for *SLFN11* (Figure 5A). *SLFN11*’s wide range of expression in SCLC samples (Figures 6E-F) and SCLC cell lines ^4^ as well as its predictive value to predict response to chemotherapeutic agents targeting DNA replication ^28,29^ (see next section) qualifies it as a quantitative biomarker. Figure 5A shows that high *SLFN11* expression in the untreated TU dataset is associated with improved survival. This observation agrees with a recent publication demonstrating preferential response of SLFN11-expressing cells to the DNA-targeted chemotherapies used to treat SCLC ^30^.

**Figure 6.**
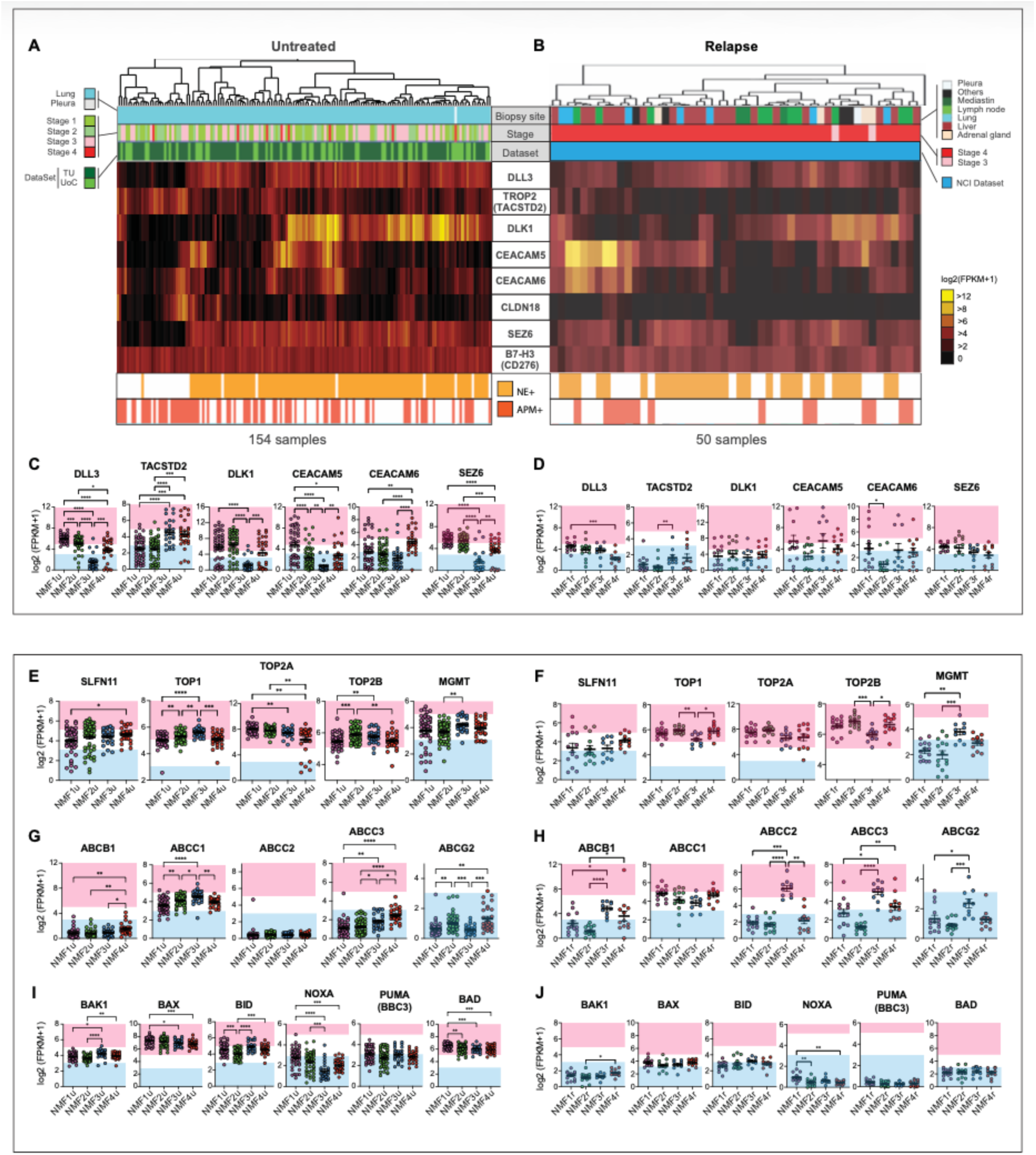
Predictive biomarkers with focus on ADCs for untreated patients (left panels A, C, E, G & I) versus patients after relapse (right panels B, D, F, H & J). **A-B**. Heatmaps showing the differential expression of 8 selected ADC targets. Samples were clustered based on the ADC target surface receptor genes listed between panels A and B. DLL3 and TROP2 (TACSTD2) are both FDA-approved targets. Expression of *DLK1*, *CEACAM5*, *CEACAM6*, *CLN18*, *SEZ6* and *B7H3* are displayed because of their selective expression in subsets of patients. The top parts of the figure show for each patient the biopsy site, clinical stage, sample type and dataset. The bottom part shows for each patient whether the sample is enriched for the neuroendocrine (NE) and antigen-presenting-machinery (APM) signatures. **C-D**. Relationship between the listed ADC antibody targets and the NMF subgroups (NMFu and NMFr stand for the NMFs of samples from untreated and relapse patients, respectively. The blue-shaded areas include samples with low gene expression (log2(FPKM+1) <3) and the pink-shaded areas samples with high expression (log2(FPKM+1) >5). **E-F**. Expression of drug targets/biomarkers in the different NMF subgroups. **G-H**. Expression of drug efflux transporters. **I-J**. Expression of major pro-apoptotic genes in the different NMF subgroups. Statistical comparisons were performed by Student t-test. **** indicates p < 0.001; ** p < 0.01; * p < 0.05.

SCLC TumorMiner also allows multivariable analyses by combining different genes or signatures. Figure 5B-C shows the predictive value of combining the expression of *SEZ6*, *SRSF1* (encoding the Serine and Arginine Rich Splicing Factor 1) ^26^ with the proliferation marker *MKI67* in the 150 untreated patients (HR 2 groups = 1.7; p-value = 0.016). In the multivariable Cox analysis, age, sex, and SEZ6 expression were independent predictors of survival, with older age, male sex, and high SEZ6 associated with increased hazard. SLFN11 showed a borderline protective effect, while tumor stage and other genes (MKI67 and SRFS1) were not significant after adjustment. These results suggest that SEZ6 provides prognostic value beyond conventional clinical factors and may improve risk stratification. Additional predictive biomarkers can be searched using the Lasso analysis option (highlighted in yellow at the bottom left in Figure 5A).

The prognostic value of each of the eight NMF gene signatures can also be explored in the “Prognosis Biomarkers” tool of SCLC TumorMiner. Figure 5D shows that at diagnosis before treatment, patients classified as NMF1u appear to have a better prognosis than patients in the MMF2u subgroup. This appears different for patients at relapse, where the NMF3r group appears at higher risk (Figure 5D).

The prognostic value of each of the eight gene signatures can also be explored in SCLC TumorMiner. Figure 5E shows the good prognosis value of the APM, immune, and Stroma signatures while male patients tend to have poor survival in the untreated datasets. This observation further indicates that tumors with high immune and stroma infiltration respond better to treatments.

None of the NAPY genes show significant predictive value, even in the untreated patients despite their broad differential expression (see Figure 1). Similarly, *MYC*, *MYCL*, *REST* and *NOTCH* do not show significant predictive value. On the other hand, consistent with a recent study reporting that high expression of *SRSF1* is linked with poor survival in SCLC patients ^26^, we observe that high expression of *SRSF1* is a risk factor for the untreated patients (Figure 5F). Single gene analyses also confirm the poor prognostic value of *SEZ6* in the untreated patient datasets and the good prognostic value of high *SFLN11* expression in the patients at relapse (Figure 5F).

### Predictive biomarkers for cell surface targets

Thirteen Antibody Drug Conjugates (ADCs) have been approved by the FDA in the past 6 years. Datroway^®^ (Datopotamab Deruxtecan), the most recent (January 2025) delivers the TOP1 inhibitor Deruxtecan to triple-negative breast cancers (TNBC) expressing TROP2 (encoded by *TACSTD2*). Notably the other TROP2-targeted TOP1 ADC, Trodelvy® (Sacituzumab Govitecan) has been granted Breakthrough Therapy Designation for ES-SCLC patients ^31^. CAR-T cells and bi- and tri-specific T-cell engagers are also in active development with the recent approval of the T-cell engager Imdelltra^®^ (Tarlatamab-dlle) for ES-SCLC ^32^. However, DLL3 or TROP2 are not validated companion diagnostic (CDx) for patient selection and, to our knowledge, RNA-seq has not been fully explored in clinical trial analyses.

Figures 6A-B display the expression of selected clinically relevant cell surface target genes (Table S9). As shown in Figures 6A and C, *DLL3* shows a broad range of expression in the untreated patient samples with the highest levels in the NE samples matching the NMF1u & NMF2u groups (Figure 6C). Notably, in the samples of patients at relapse, DLL3 expression is at least 3-fold lower and lacks clear discrimination among the NMFs groups (compare Figures 6C and D), consistent with the reduced *ASCL1* expression at relapse (see Figure 1A). These results suggest the potential of measuring *DLL3* transcripts as a CDx in front-line SCLC.

TROP2 (encoded by *TACSTD2*) appears generally low at relapse (Figure 6B) while, in the untreated patients, its expression is high in samples with low NE score and high APM score (Figure 6A). Accordingly, TROP2 expression matches the NMF3u (*POU2F3*-endriched) and NMF4u (Immune-enriched) clusters (Figure 6).

Figure 6 (upper panels A-D) displays the expression of additional cell surface targets for ADCs that are in clinical trials (Table S9). *DLK1* (Delta Like Non-Canonical Notch Ligand 1) is highly expressed untreated samples with NE signature (Figure 6A) and less in relapse tumors (Figure 6A-D). NMF analyses show high *DLK1* expression in the NMF1u and NMF2u and in approximately 50% of the NMF4u group (Inflammatory and Immune) (Figure 6C). These results support the ongoing clinical trial testing a DLK1-ADC in NE tumors ^33^.

Two of the carcinogenic embryonic antigens (encoded by *CEACAM5* and *CEACAM6*), which are targeted with ADCs in clinical trials for colon, gastric cancers and NSCLC (Table S9), are highly expressed in the samples classified as NMF1u and NMF2r (Figures 6A-D). These results suggest the potential of CEACAM5/6-targeted ADCs in SCLC and the potential of measuring CEA in the blood of SCLC patients as a CDx biomarker ^34,35^.

*CLDN18* (coding for Claudin 18), an ADC target for pancreatic cancer (Table S9) is highly expressed in a few untreated patients; yet, not in relapse patients (Figures 6A-B). As discussed above, *SEZ6*, a downstream target of ASCL1 ^36^ is highly expressed across untreated and relapse samples matching the NE signature with only partial overlap with *DLL3* expression (Figures 6A-D). Finally, B7H3 (encoded by *CD276* and belonging to the immunoglobulin superfamily) is expressed in a broad range of samples (Figures 6A-D). Together, these results validate the potential of DLL3-, SEZ6-, DLK1-, CEACAM-, CLDN18 and B7H3-targeted ADCs in SCLC ^37^.

### Predictive pharmacodynamic (PD) biomarkers related to the ADC payloads

The bottom panels of Figures 6 summarize the expression of known intracellular drug targets (Figure 6E-F), drug efflux transporters (Figure 6G-H) and proapoptotic genes (Figure 6I-J).

*SLFN11* (Schlafen 11) shows a broad expression range (over 8-fold) in both the untreated and relapse patient samples, in agreement with SLFN11’s wide distribution across the SCLC cell lines irrespective of the NAPY subgroups ^4^ (https://discover.nci.nih.gov/SclcCellMinerCDB/). Consistent with *SLFN11* being an immune- and interferon-response gene ^28^, the NMF4 inflammatory/immune subgroups tend to show highest *SLFN11* expression both in the untreated and relapse patient samples (Figure 6E and F). The broad range of SLFN11 expression suggests that further studies are warranted to confirm the relevance of SLFN11 to predict response of patients treated with lurbinectedin, topotecan and TOP1 inhibitor payload ADCs ^38–42^.

Since TOP1 is the primary cellular target of topotecan, irinotecan and TOP1-ADCs, and TOP2 the target of etoposide, we looked at *TOP1* and *TOP2* expression (Figure 6E-F). Both are highly expressed in the samples from untreated and relapse patients as expected for housekeeping genes. Thus, *TOP1* and *TOP2* levels are irrelevant as predictive biomarkers.

Lack of O-6-Methylguanine-DNA Methyl-Transferase (MGMT) predicts the activity of DNA alkylating agents, and foremost of temozolomide ^43^, which is occasionally used in SCLC ^44^. Figure 6 (right panels E & F) shows low expression MGMT in some samples, suggesting the potential of testing *MGMT* expression for temozolomide treatments.

Drug efflux transporters of the ABC family are a major cause of resistance to tubulin inhibitors (vinca and auristatin derivatives) that are commonly used as ADC payloads. Yet, measuring ABC transporters is not done in clinical routine. Figure 6G-H displays the expression of the main ABC transporter genes. *ABCB1*, which encodes P-glycoprotein (PgP) has notably low expression both in the untreated and relapse samples, while *ABCC2* and *ABCC3* show significant overexpression in the NMF3r cluster (Figure 6H). These results suggest that measuring ABC transporters should be considered for ADCs with anti-tubulin payloads.

Because broad resistance to chemotherapy is dependent on reduced apoptosis, we explored apoptotic genes. Figures 6I-J and S9C show global suppression of proapoptotic gene expression in relapse patient, which may account, in part for the overall resistance of relapse patients to chemotherapy.

### Analysis of individual patients using the “MyPatient” tab of SCLC TumorMiner

Going forward, we wish to propose SCLC TumorMiner as a prototype clinical interface for classifying tumors and for predicting risk factors and rational therapies (Figure 7A). This feature is included in the “My patient” tab of SCLC TumorMiner (Annotation #7 at the top of Figure 1A).

**Figure 7.**
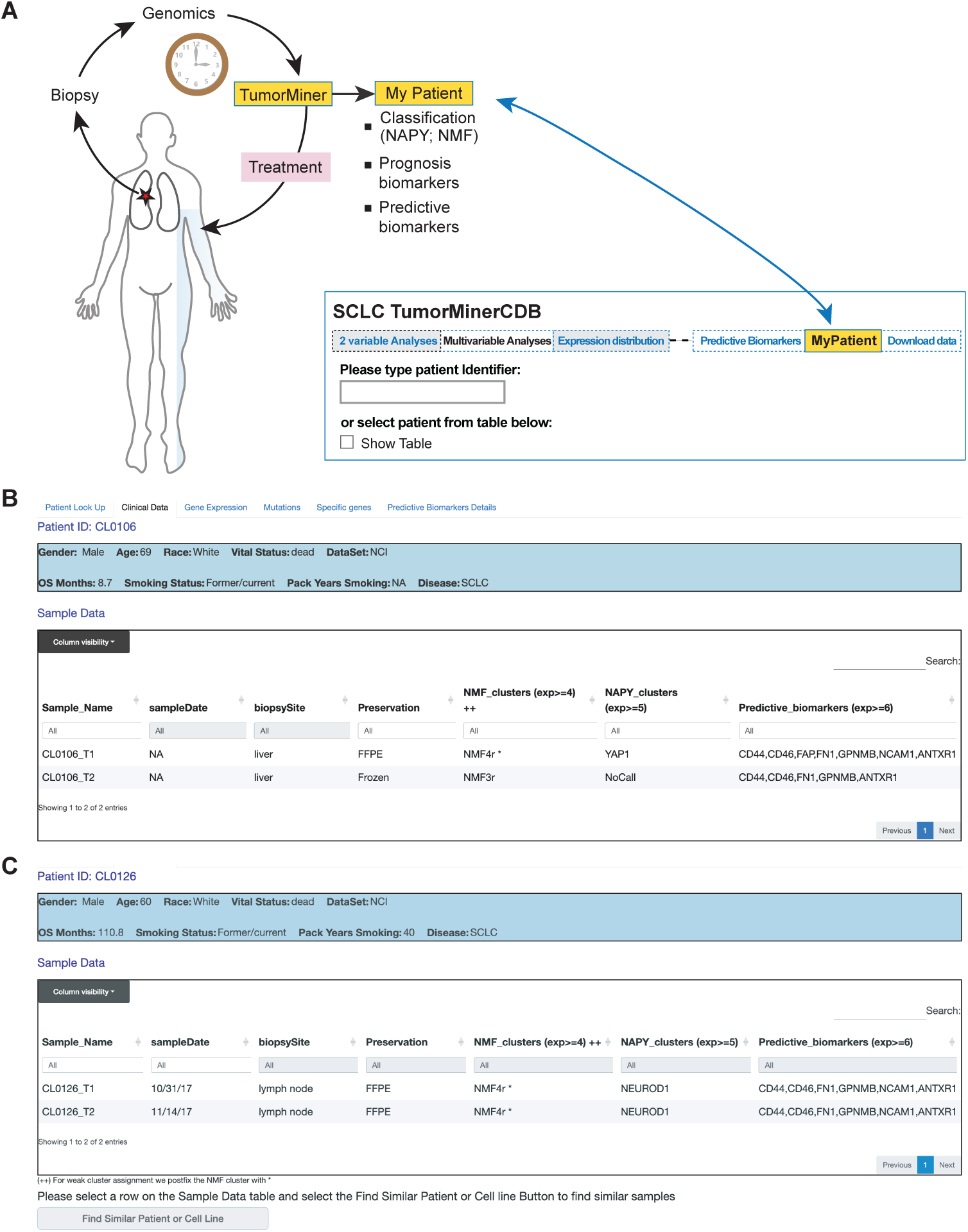
MyPatient module of SCLC TumorMiner. **A**. Proposed use of the “MyPatient” module. Genomic analyses are performed immediately after biopsy and uploaded into Tumor_Miner to provide results guiding the patient treatment as genomic classification, prognostic and predictive biomarkers are automatically displayed using “MyPatient” tab. **B**. Snapshot of the “MyPatient” tab in SCLC TumorMiner. Users are prompted to enter a “patient identifier” or select a patient from the available table. Here the patient has two samples from liver biopsies: one frozen and the other from FFPE. **C**. Snapshot of the “MyPatient” tab for patient “CL0126” who has two lymph node biopsy samples taken at 2-week interval.

As discussed above (see Figure 1B), all the untreated patient samples can be assigned to at least one of the NAPY groups based on the high expression cut-off of log2(FPKM+1) > 5 defined above. Therefore, we have included the NAPY classification to effectively characterize the samples from untreated patients in the “My patient” tab of SCLC TumorMiner. The situation is different for the relapse SCLC samples. Indeed, based on the expression cut-off of log2(FPKM+1) > 5, less than half of the samples (44%) can be assigned to the NAPY groups, with the YAP1 group (SCLC-Y) being prominent (22%) (see Figure 1B).

Faced with the limitation of the NAPY classification for the relapse patient samples and the positive discriminatory value of the NMF classification (see Figures 3 and S4), we developed an algorithm to classify the relapse samples based on NMF. Figure S7 shows the average expression scores for the NMFr clusters of the 50 samples from the NCI relapse patients. Based on this plot, we chose a threshold of 4 to assign samples to each NMFr cluster. This approach enables the computation of mean average NMF scores for individual samples. On the website, we implemented this approach to score automatically individual samples in the “MyPatient” tab of SCLC TumorMiner. Upon typing a patient identifier (code) (Figure 7B), SCLC-TumorMiner displays both NAPY and NMF classifiers for cataloging individual patient samples (Figure 7B-C).

The tab “My Patient” also displays predictive biomarkers for therapies (Figure 7B-C). This is done by the TumorMiner software, which queries a dataset listing 63 cell surface targets for ADCs currently FDA-approved and in clinical development (Table S9). Our website displays the cell surface targets expressed above a threshold of 6 (log2(FPKM+1)). For example, in Figure 7C, the displayed patient tumor sample overexpresses NCAM1 and FN1 implying that the patient could be candidate for ADCs comprising NCAM1 or FN1 antibodies.

These results highlight the translational potential of the SCLC TumorMiner platform for individualized patient care. While traditional NAPY classification remains useful for untreated tumors, its limited applicability to relapsed samples underscores the importance of alternative classifiers. By implementing NMF-based transcriptomic classifiers and integrating biomarker-based predictions for targeted therapies, the “MyPatient” module can provide real-time, personalized tumor profiling. Our goal is to facilitate not only tumor subtype identification but also actionable therapeutics, bridging molecular characterization with clinical decision-making in SCLC. As the platform evolves with more comprehensive datasets, it holds promise for guiding precision medicine strategies in both newly diagnosed and treatment-refractory patients.

## DISCUSSION

Mining cancer patient clinical and genomics data is critical for understanding a disease, discovering biomarkers, predicting outcomes, optimizing treatments, and ultimately shifting patient care to a logic-based and flexible paradigm. During the last decade, new tools have emerged such as cBioPortal (http://cbioportal.org) ^45^ or Xena (https://xena.ucsc.edu) ^46^, GEPIA2 (http://gepia.cancer-pku.cn/) ^47^, UALCAN (https://ualcan.path.uab.edu/) ^48^ or Survsig (https://survsig.hcemm.eu/), offering different functions for genomic data visualization and analyses.

Here we introduce “SCLC TumorMiner”, a web application for SCLC patient genomics. Its architecture and displays are constructed like SCLC-CellMinerCDB (https://discover.nci.nih.gov/SclcCellMinerCDB), which enables users to compare tumor samples with a broad collection of SCLC cell lines ^4,49^. SCLC TumorMiner complements existing tools for mining SCLC patient clinical and genomics information, such as cBioPortal, Xena or Survsig. It is also meant to improve access and interpretation for physicians and their patients. SCLC TumorMiner has its own specific datasets and unique functionalities to build predictive models and generate prognostic gene sets. It is the first tool to enable comparison between a patient sample and other patient samples or cell lines. Table S10 showcases the distinctive values of SCLC TumorMiner with other tools.

SCLC TumorMiner (https://discover.nci.nih.gov/SclcTumorMinerCDB/) has a battery of unique tools (see Figure 1A) allowing real-time display of omics data for individual genes and individual patient samples (*2-Variable Analyses*, *Prognosis Biomarkers* and *Predictive Biomarkers*) or for multiple genes to explore molecular networks (*Multivariable Analyses* and *Compare Patterns* tools) (see Figure 1A). It integrates 8 pathway *Signatures* relevant to SCLC biology (NE, RepStress, CES, APM, Immune, EMT, Stroma, Y-Chrom), 50 Hallmark signatures and 63 cell surface targets relevant to the current development of Antibody Drug Conjugates (ADCs), Bispecific T-cell engagers (BiTEs) and CAR-T cell therapies. Data can be downloaded from the SCLC TumorMiner website for further analyses, as demonstrated in this report by the different plots comparing gene expression in untreated and treated patients and in samples from different NMF groups.

A biologically relevant feature of TumorMiner is its ability to explore the logic of molecular networks/pathways based on the premise that individual components of molecular pathways are quantitatively linked and significantly correlated ^4^. Figures 2 and S2 demonstrate that each of the NAPY genes has its own network (genomic identity) across tumor samples and that these networks are highly conserved across patient samples and cell line models (Figure S2C), implying their robustness. A notable finding is the interconnection between the Notch, the YAP/TAZ and the ASCL1-NE pathways whereby expression of the YAP/TAZ pathway is tightly linked to activation of the Notch pathway; and vice-versa that activation of the NE (ASCL1-SEZ6) pathway is associated with expression of the Notch ligands (DLL3, DLK1, DLL1, DLL4 and JAG2) and negatively correlated with the Notch receptors and their transcription effector REST (see Figure 2F). These results suggest feedback loops involving the non-NE pathways including YAP/TAZ and the Notch receptors, and the NE pathway including ASCL1 and the Notch ligands ^21,22^. Whether and how YAP/TAZ functions upstream of the Notch transcription factor REST remains to be established.

Cancer cell plasticity and heterogeneity are two outstanding characteristics of SCLC ^50–56^. This is exemplified by the fact that at relapse, patients become resistant to chemotherapy after initially responding generally dramatically. Accordingly, SCLC TumorMiner reveals key genomic differences between the samples of untreated and relapsing patients. One such difference is the loss of the bimodal expression of NEUROD1, ASCL1 and POU2F3 and the upregulation of YAP1 at relapse (see Figure 1). This could have, at least two not mutually exclusive explanations: 1/ differentiated cancer cells expressing ASCL1 (NE cells) or POU2F3 (Tuft-ionocyte-like cells) are eliminated by the 1^st^ line of chemotherapy (see Figures 1A and S1B), leaving behind less differentiated resistant cells; 2/ chemotherapy is less effective toward the less differentiated and more plastic tumor cells with basal cell characteristics ^55^, which remain in the tumor and become prominent at relapse.

Another notable difference between untreated tumors and tumors at relapse is the significant downregulation of MYCL at relapse (see Figure 2G). This reduction is likely consequential to the fact that ASCL1 is a transcriptional driver of MYCL ^57^ (as visualized in Figures 2B) ^57^ and that ASCL1 expression is significantly suppressed in relapsed tumors (see Figures 1A and B).

With respect to the chemoresistance of relapsed tumors, it is likely multifactorial. Indeed, while the expression of SLFN11 and some of the proapoptotic genes (BAD, BAX, BID, NOXA and PUMA) is high in chemotherapy-naïve patient samples, it is markedly reduced in relapsed tumors (see Figures 6), suggesting that the initial response of patients to various chemotherapies is driven, in part by pro-apoptotic genes and SLFN11. Conversely, resistance may involve downregulation of SLNF11 and the cell death pathways. Whether these pathways are epigenetically suppressed in relapse tumors and could be reactivated by HDAC and EZH2 inhibitors remains to be tested ^10,39,58^.

Bispecific antibodies T-cell engagers (BiTEs), antibody drug conjugates (ADCs), and CAR-T are in full development. Regarding the therapies targeting DLL3, Tarlatamab (Imdelltra^®^) and Zocilurtatug Pelitecan (Zl-1310) (NCT06179069), expression of DLL3 is generally lower at relapse than in untreated patients, consistent with DLL3 being a direct transcriptional target of ASCL1, which is significantly suppressed at relapse (Figures 6 & S9) ^4,57^. This may justify moving Tarlatamab and Z-1310 to 1^st^ line therapy. Also, since DLL3 IHC analyses have until now failed to predict responses ^59^, using RNAseq could identify responders (approximately 40%) to Tarlatamab ^32,59^ and Z-1310 (NCT06179069), thereby avoiding unnecessary toxicity and cost. Surprisingly, expression of TROP2 (encoded by *TACSTD2*) is generally low both in the untreated and relapse patient samples. This questions the potential of Trodelvy^®^ (Sacituzumab Govitecan) or Datroway^®^ (Datopotamab deruxtecan) in SCLC patients irrespective of their disease stage.

Based on our observations, SEZ6 appears promising for two of the ADCs in clinical development: ABBV-011 and ABBV-706 (see Table S9). Yet, like DLL3, SEZ6 is a downstream transcriptional target of ASCL1 ^36,37^ (see Figure 2) and is therefore likely to be more effectively targeted in untreated patients. By contrast, B7H3 (encoded by *CD276*), the target of two ADCs in advanced clinical trial for ES-SCLC: Risvutatug Rezetecan (GSK5764227) and Ifinatamab Deruxtecan (DS-7300) (Table S9) appears highly expressed in a significant fraction of samples from relapsed patients in SCLC TumorMiner. Our systematic screening for other cell surface targets reveals that CEA (carcinoembryonic antigen), which is encoded by *CEACAM5* and *CEACAM6*, is highly expressed in multiple samples from relapse patients (Figure 6), suggesting the potential of CEA-ADCs ^60–62^. An advantage of targeting CEA is that, in addition to measuring CEACAAM5/6 transcripts and by IHC in tumor samples, clinical trials could include CEA blood samples ^63,64^. We also found that TRPM5, which encodes a sodium channel transporter, is selectively expressed in the POU2F3-expressing tumors and that GFRA1 (Glial Cell Line-Derived Neurotrophic Factor Receptor Alpha) and SEMA6A (encoding one of the semaphorins) ^14^ are specifically expressed in NEUROD1 expressing SCLC (Figure 2), suggesting they might be considered as novel cell surface targets for SCLC. These analyses demonstrate the potential use of SCLC TumorMiner for discovering novel cell surface therapeutic targets for ADCs, T-Cell Engagers (BiTEs) and CAR-T.

As we move toward precision medicine, multivariable analyses will be needed. For instance, for a given cell surface target, clinicians may have to choose between an ADC with a TOP1 or a tubulin payload or a T-cell engager (BiTE). To illustrate this point, one could propose guiding payload choice with at least two biomarkers: SLFN11 and ABC drug efflux transporters. If the tumor expresses high SLFN11 and drug efflux transporters, TOP1 inhibitors will likely be superior to a tubulin inhibitor ADC, and conversely a tubulin inhibitor ADC payload would be logical for tumors with low SLFN11 and ABC transporters expression. For immune checkpoint inhibitors and BiTEs, one could consider testing the predictive value of high APM and Immune signatures, which would reflect the presence of immune cells in the tumor, or tumor intrinsic features such as high APM or NOTCH1 expression ^65,66^.

Ultimately, we speculate that “*MyPatient*” could be developed for clinical practice as bulk RNA-seq and DNA-seq can routinely be performed. Sequencing results and clinical parameters could be loaded into SCLC TumorMiner. Additionally, the Tumor_Miner architecture and interface could be readily expanded to integrate protein expression (proteomics) and epigenetic (DNA methylome) parameters as we have already done in our CellMiner websites (https://discover.nci.nih.gov/). “My_Patient” could provide a medical interface displaying a patient’s genomic parameters, dictating which therapy would be most appropriate and predicting risk-factors. Finally, one could anticipate future integration of natural-language query interfaces or Chatbot to enhance user accessibility.

## LIMITATIONS OF THE STUDY

A limitation of the current study is the technical and clinical heterogeneity of the cohorts. RNA-seq data were generated from both fresh frozen and FFPE samples, with untreated cohorts primarily derived from frozen tissues and the relapse NCI cohort including both frozen and FFPE samples. Analysis of the NCI samples, including 8 frozen biopsy samples and 42 FFPE samples, shows that preservation methods only explain 2.9% of the variance, which is quite low, especially compared to the biopsy site, which explains 10.8% of the variance. Hierarchical clustering analysis of the 204 samples from the 3 datasets (UoC, TU and NCI) also shows a clear separation between NCI (relapse) and the other samples (Untreated). The NCI relapse samples are mostly FFPE, whereas the untreated samples are all Frozen. We cannot remove the effect of the preservation because it is confounded with sample type (relapse vs. untreated). Thus, no batch effect was applied for all 3 datasets, and we believe that bulk RNA-seq of patient samples can give a valuable estimate of gene expression for highly expressed genes [log2(FPKM+1) > 5]. Adding more samples will increase the statistical power and robustness of SCLC TumorMiner. However, a challenge for cross-database analyses is to access large and well-annotated datasets. Material transfer agreements commonly restrict the sharing of FASTQ files and associated clinical data across institutions. **“**MyPatient” module only provides a prototype, and users will have the option to integrate our TumorMiner platform into Electronic Medical Record (EMR) systems to provide “real-time” evaluation of tumor Omics for precision medicine.

## Author Contributions

Y.P. supervised the study. Y.P., F.E., Y.A., devised the concept. Y.A., D.T., A.T., A.L., J.D.R, F.A., K.A. and M.R. data collection and clinical annotations. Y.P., F.E., A.D., S.V., D.T., Y.A., A.L., R.S., N.R., A.T., C.R., A.T. and W.R performed data analysis. F.E., J.W. and A.Te. built the website. Y.P., F.E., A.D., A.L., A.T., M.A., N.R., C.R., W.R wrote and critically reviewed the manuscript. All authors reviewed the results and approved the final version of the manuscript.

## Data Availability Statement

The clinical information for all four datasets provided in Supplementary Data and normalized FPKM expression data is available at https://discover.nci.nih.gov/SclcTumorMinerCDB/.

## Acknowledgments

Our work is supported by Center for Cancer Research, National Library of Medicine and Developmental Therapeutics Branch, National Cancer Institute. We thank the NCI CCR Collaborative Bioinformatics resource (CCBR) team for their help in running the RNA-seq and Exome-seq pipelines.

## STAR METHODS

### RESOURCE AVAILABILITY

**Lead Contact:** Further information and requests for reagents may be directed and will be fulfilled by Lead Contact Yves Pommier

**Material Availability**: this study did not generate new unique reagents

## Data and code Availability

- **Data:** the data used for the analysis can be downloaded from the website at https://discover.nci.nih.gov/SclcTumorMinerCDB/ or obtained from Zenodo at https://doi.org/10.5281/zenodo.18343141
- **Code:** the website code is publicly available in GitHub at https://github.com/cbiit/SclcTumorMinerCDB
- **Other:** Any additional information or script required to reanalyze the data reported in this paper is available from the lead contact upon request

## EXPERIMENTAL MODEL AND STUDY PARTICIPANT DETAILS

Patient clinical and genomics data were collected from our DTB clinic at the Center for Cancer Research, NCI-NIH, the University of Cologne (UoC) and Tongji university (TU). Our genomics patient data currently includes a total of 204 samples from 189 patients with RNA-seq and/or whole-exome sequencing (WES) samples. The NIH samples were obtained from patients enrolled in IRB-approved protocols and had consented to the analysis of their tumors. Clinical data from NIH and collaborators were gathered and manually curated by a team of DTB physicians. They integrate patient demographics, survival, treatment with response, and sample information, including sample type, biopsy site, tumor stage and pathology indicators (see Table S1). NCI patient samples were correlated with treatment response, and for each patient we selected the first drug(s) or treatment(s) post-sampling as well as its response class (CR: complete response, PR: partial response, PD: progressive disease or SD: stable disease). Due to the current limited number of treatments, we decided that the response will be of value 1 (good responder: PR, CR) or 0 (bad responder: SD, PD).

## METHOD DETAILS

### Genomic content and Infrastructure

All genomics patient data were processed with our on-site data-handling pipelines using the raw data. We generated 204 RNA-seq and 114 WES normalized data. Treatment responses were available for 14 patients. Table S11 shows details by disease, data sets and experiments.

We also computed for each sample gene expression Hallmark enrichment and genomics signatures scores such as the NE, RepStress, EMT, APM, the level of stroma and immune infiltrate contamination or a new Y chromosome signature score. Table S12 presents the list of signatures as well as the phenotypic data queryable on SCLC TumorMiner.

SCLC TumorMiner has similar design as the CellMinerCDB tool and websites (https://discover.nci.nih.gov/) dedicated to mining pharmaco-genomics data across broad ranges of cancer cell line sets and databases. SCLC TumorMiner is implemented as a shiny application and hosted using Posit Connect on CBIIT servers at https://discover.nci.nih.gov/SclcTumorMinerCDB/.

### RNA sequencing and batch assessment

The NIH/NCI (relapse) cohort was composed predominantly of FFPE specimens (n=42), with a limited number of frozen samples (n=8) using the Illumina access protocol, whereas the Universities of Cologne (UoC) and Tongji university (TU) were from Frozen using PolyA protocol. All the SCLC samples were processed by running the NCI CCBR RNAseq pipeline (https://github.com/skchronicles/RNA-seek.git). STAR (2.7.6a) was run to map reads to hg38 reference genome (release 36). Then RSEM was used to have gene expression values in log2(FPKM+1). For the NCI dataset, we checked whether the way the samples were preserved—Frozen or FFPE—affects gene expression, along with other factors like biopsy site, sex, and age. Analysis of the NCI samples including 8 frozen biopsy samples and 42 FFPE samples shows that preservation methods only explain 2.9% of the variance, which is quite low, especially compared to the biopsy site, which explains 10.8%. All untreated samples are frozen. Because the preservation method is confounded with sample type (relapse vs. untreated), its effect cannot be disentangled (See Figure S8). Thus, no batch effect was applied for all 3 datasets.

### Mutations

Somatic mutations were based on paired tumor-normal samples and ran with the CCBR exome sequencing pipeline (https://github.com/mtandon09/CCBR_GATK4_Exome_Seq_Pipeline). Briefly, BWA MEM (version 0.7.17) was run to map reads to the hg38 reference genome. Then Mutect2 in GATK 4.2 was used to call the variants.

Across 98 Tongji University paired normal and tumor exomes, the median of the mean target depth was 161.57× (IQR 139.32–189.27×), and a median of 97.1% (IQR 96.2–97.7%) of target bases were covered at ≥30×. For NCI, across 16 paired normal and tumor exomes, the median of the mean target depth was 97.75× (IQR 62.9–143.5×), and a median of 90.9% (IQR 79.9–95.42%) of target bases were covered at ≥30×. Sample coverage details are provided in supplementary Tables S13 and S14.

Variants with 6 or more total filtered reads were selected for display. We used Annovar (hg38 version) to annotate the mutations, including their predicted effect on amino-acid sequence and whether they are predicted to be deleterious, as well as their prevalence in the ExAC normal population. A mutation was called deleterious if its predicted effect on the RNA sequence was frameshift, nonsense, splice-site, or a missense mutation with a SIFT score <0.05 or PolyPhen2 HDIV score>=0.85. We selected all deleterious variants for the mutation gene-level score.

The fraction of alternate alleles (the mutation ratio) was used as the probability that any cell has that mutation. Furthermore, we assumed that the probability that any mutation is present in a gene is independent of the probability of any other mutation on that gene. Given those assumptions, we calculate the overall mutation probability for that gene in any cell as:

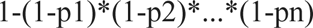

where p1, p2, …, pn are the mutation ratios for each individual mutation on that gene. 1-pj is the probability that jth mutation is not present for a particular cell. Thus, the product is the probability that none of the mutations are present and 1 minus the product is the probability that at least one mutation is present. The expression gives a number that is between 0 and 1, which we multiply by 100 to get the mutation score for that gene.

### Tumor Mutation Burden (TMB)

The TMB was computed using the R package MAFtools ^67^ based on Mutect2 somatic variants that have variant allele frequency (VAF) >= 5%.

### Genomic data for survival

If a patient has multiple samples, we select the primary tumor sample (if available) for survival analysis or we choose one based on biopsy site. The list of selected samples is downloadable from the website.

## QUANTIFICATION AND STATISTICAL ANALYSIS

### Unsupervised Clustering Approach

We implemented unsupervised clustering based on Non-Negative Matrix Factorization (NMF) algorithm using log-transformed normalized RNA-seq data to classify SCLC patients into molecular subtypes. NMF uses dimensionality reduction approach in which it decomposes the gene expression matrix X (n × m), into the product of two lower-dimension non-negative matrices (See Equation below), where n represents number of patients and m is number of genes.

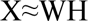

with X = n × m matrix, W = n × k matrix, and H = m × k, where k is a latent factor; optimal number of factors (k) was determined based on matrix stability using cophenetic correlation coefficient. We selected only protein-coding genes for both untreated and relapse SCLC samples, then we selected the top 5000 highly variable genes for both datasets using a non-parametric approach, Median Absolute Deviation (MAD) using “mad” function in R (version 4.2.3). After which, we performed consensus NMF clustering evaluating values of k ranging from 2 to 8, based on cophenetic correlation coefficient. k = 4 provided most stable and robust clusters for both untreated and relapse SCLC datasets (See Figure S4). Clustering was performed using “NMF” library in R (version 4.2.3).

### Pairwise Differential Expression Analysis

After defining the NMF subgroups, differential gene analyses were performed using the “Limma” package in R (version 4.2.3). To determine the up or down-regulated genes across the NMF-subgroups, we performed pairwise comparisons for untreated and relapse SCLC separately. This included all possible combinations of comparisons: NMF1 vs NMF2, NMF1 vs NMF3, NMF3 vs NMF4, NMF2 vs NMF3, NMF2 vs NMF4, NMF3 vs NMF4. In addition, each subgroup was compared with the aggregated sum of other subgroups, for instance: NMF1 vs Others (NMF2+NMF3+NMF4), NMF2 vs Others (NMF1+NMF3+NMF4), NMF3 vs Others (NMF1+NMF2+NMF4) and NMF4 vs Others (NMF1+NMF2+NMF3).

Subsequently, we utilized these pairwise comparisons to identify subgroup-specific genes for each NMF cluster for both untreated and relapse samples. Significantly upregulated genes in NMF1 in comparison to NMF2, NMF3 and NMF4 with log_2_FC >1 and p-value < 0.05 are listed as NMF1-specific genes. Similarly, this criterion was implemented for the selection of specific genes for the NMF2, NMF3 and NMF4 subgroups. The subgroup-specific genes identified using this approach were used to construct NMF-derived gene expression signatures.

### Gene Set Enrichment Analysis (GSEA)

To investigate the functional relevance of genes we performed pre-ranked Gene Set Enrichment Analysis (GSEA). From previous analysis, we obtained pairwise comparisons for each NMF groups and used t-statistic parameter, which provided magnitude and direction of differential expression to rank the genes. Ranked genes were then analyzed using the fGSEA R package (version 4.2.3) with REACTOME gene sets [https://www.gsea-msigdb.org/gsea/msigdb/human/collections.jsp#C5]. fGSEA is a fast implementation of the GSEA algorithm providing normalized enrichment scores (NES) and adjusting for multiple testing using permutation-based p-values and false discovery rate (FDR).

### Mypatient NMF cluster assignment

For each patient sample, we calculated the average expression of the four NMF signatures (see Figure S7). The assigned NMF cluster corresponds to the signature with the highest average expression, provided that this value is at least 4 (log2(FPKM+1)); otherwise, no cluster is assigned. If the highest average expression differs from the second highest by less than 1, the cluster assignment is considered weak and is labeled accordingly.

### Statistical Methods

Correlations, scatter plots, heatmaps, survival analysis, regression model predictions, and pathway enrichment scores were generated using the R-Project for statistical computing based mostly on the R packages: ggplot2, plotly, heatmaply, survival, glmnet, ggforest, GSVA and GSEABase. These packages are downloadable from the comprehensive R archive network (CRAN) or Bioconductor websites.

## KEY RESOURCES TABLE

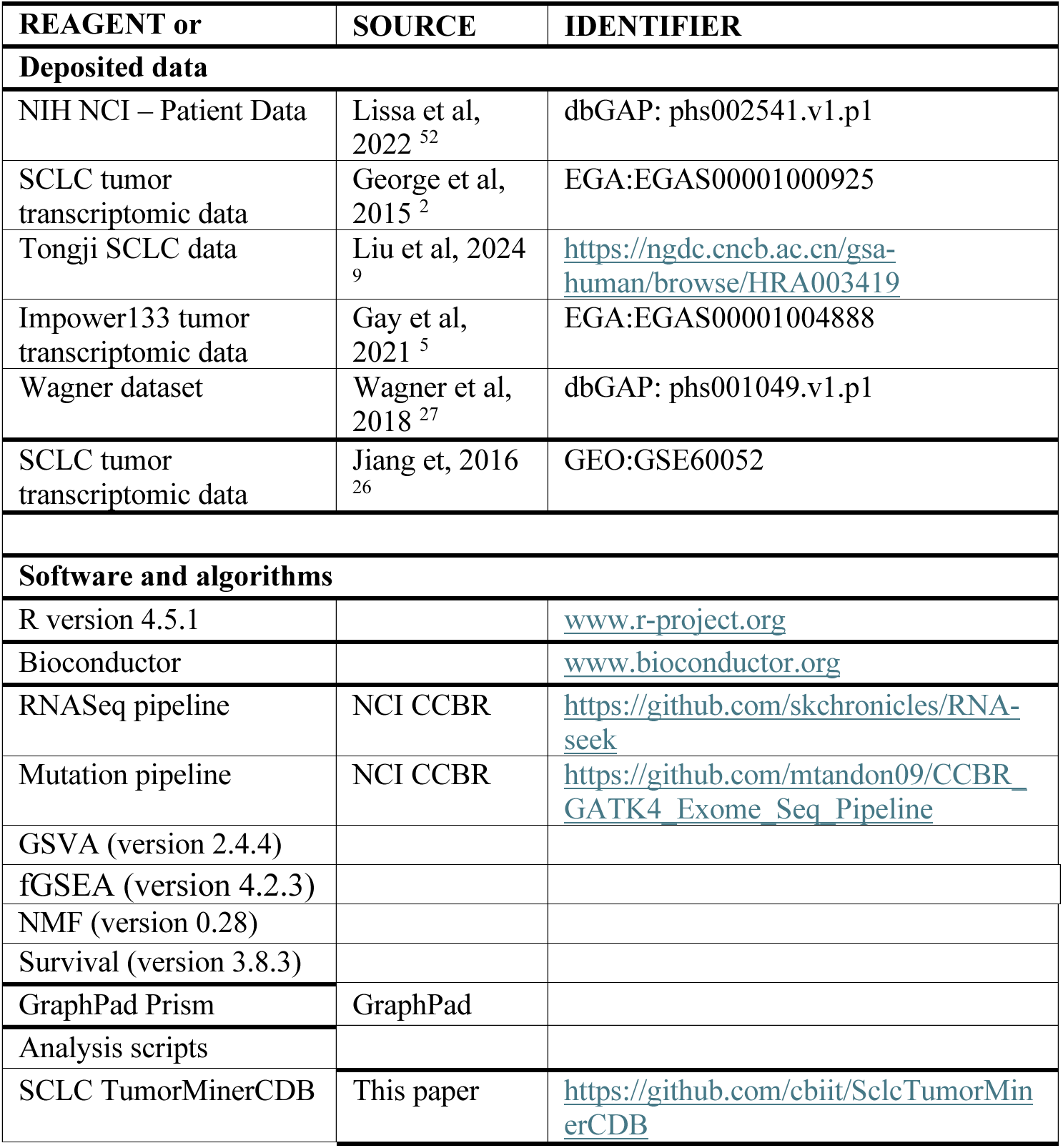

## Supplemental Figures

**Figure S1 (related to Fig 1).**
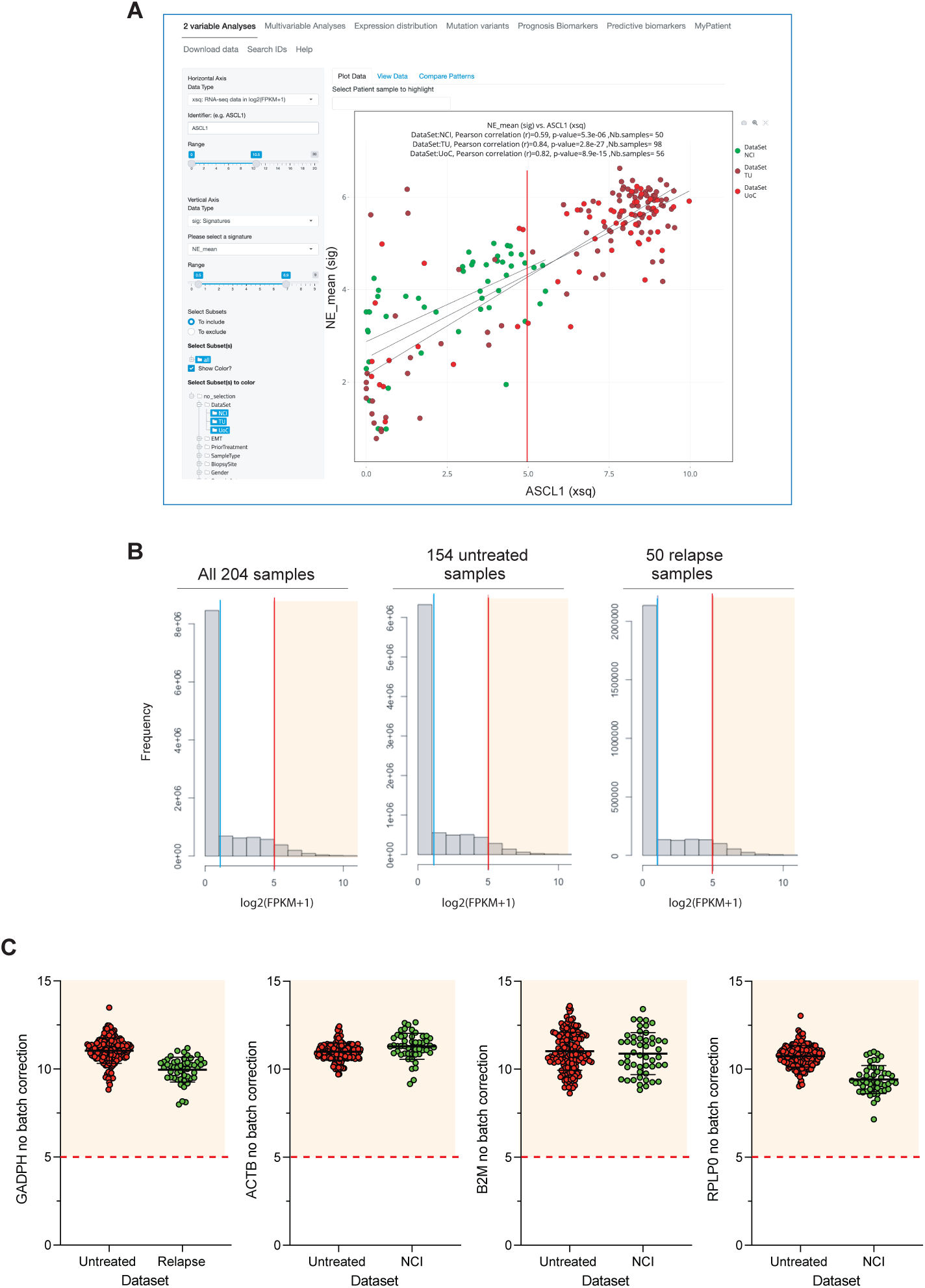
**A**. Example of a “2 Variable Analysis” snapshot of the websites displaying the correlation between *ASCL1* expression and the NE signature. **B**. Frequency distribution of gene expression across the datasets. The red line indicates the >90% cutoff (log2(FPKM+1)>5) for high gene expression. **C**. Comparative expression of 4 canonical housekeeping genes in the untreated and relapse samples. Colored areas define samples with significant expression ((log2(FPKM+1) > 5.0 in pink). Data were downloaded from the SCLC TumorMiner website using the download tabs.

**Figure S2 (related to Figs 2).**
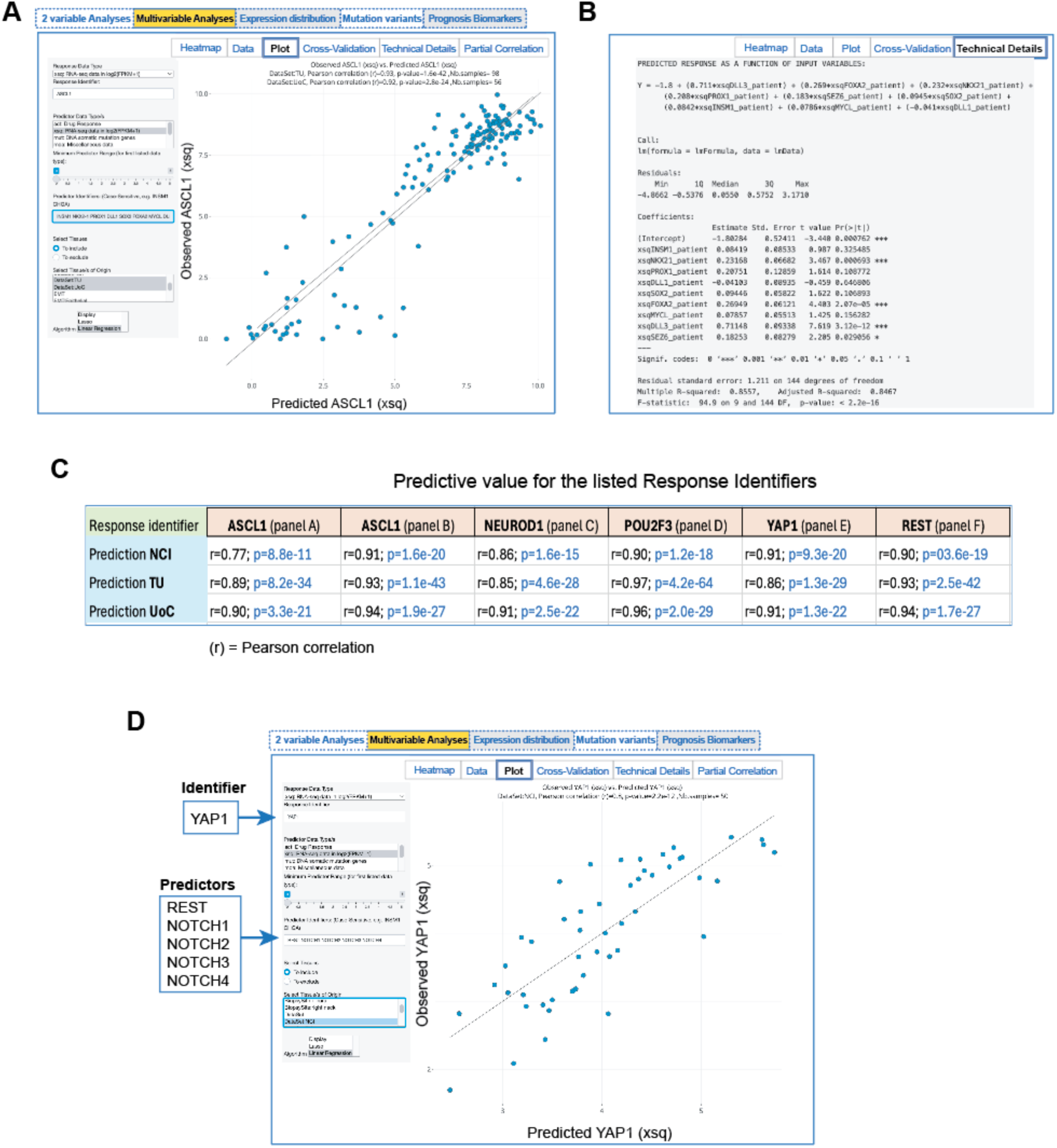
**A**. Snapshot of the plot corresponding to the predictive value of the 8 variables (“Predictor Identifiers”) listed in Figure 2A under “Multivariable Analyses”. **B**. Snapshot of the “Technical Details” corresponding to the plot shown in panel A and the Heatmap shown in Figure 2A. **C**. Linear regression analyses for the Heatmaps shown in panels A-F of Figure 2. Pearson correlation coefficients and p-values were obtained from the “Plot” tab (as shown in panel A for Figure 2A). **D**. Expression of the Notch pathway canonical genes *REST*, *NOTCH1*, *NOTCH2*, *NOTCH3* & *NOTCH4* predict the expression of *YAP1*.

**Figure S3 (related to Figure 1).**
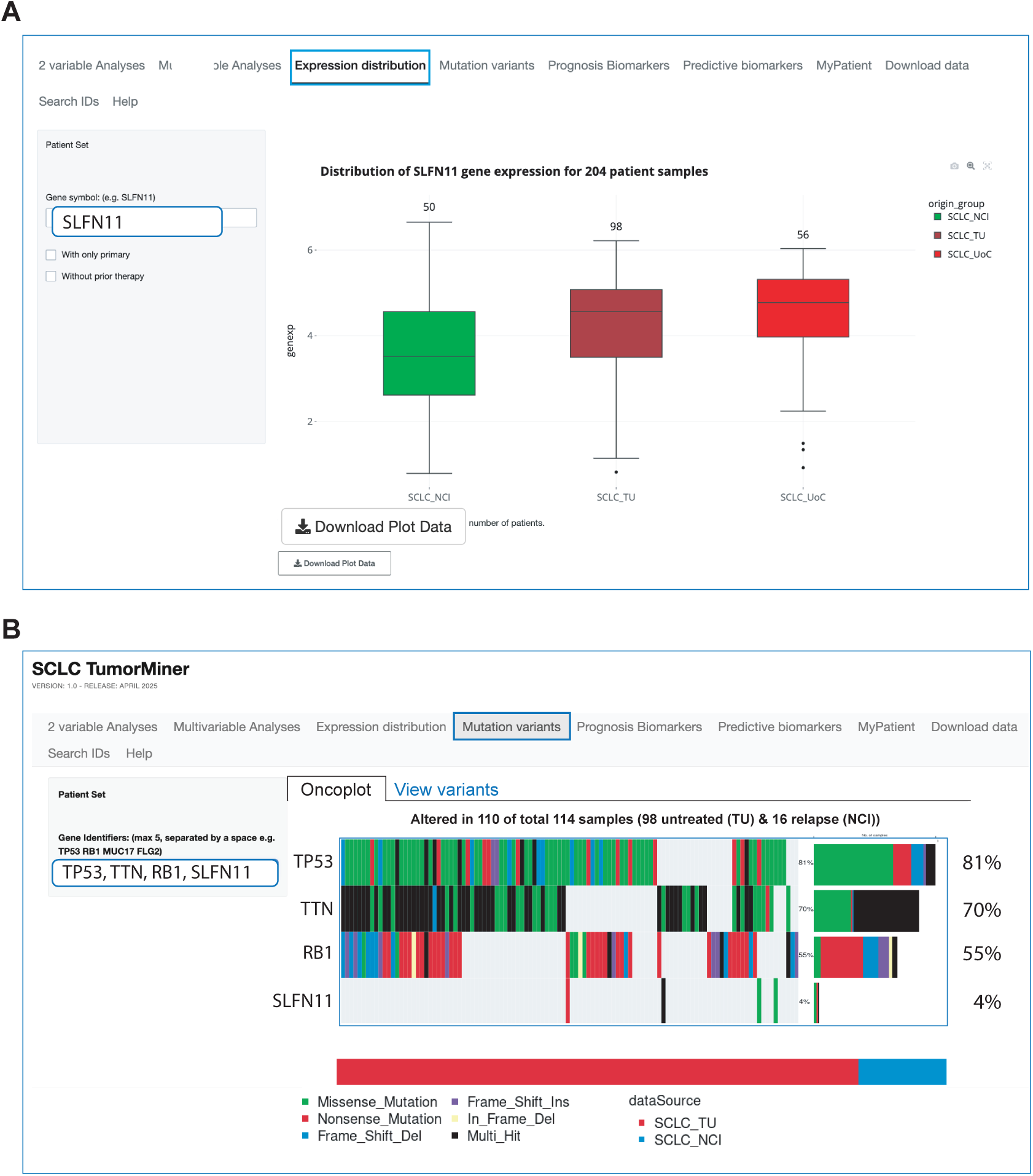
**A**. Snapshot of the “Expression distribution” tab from the SCLC TumorMiner website in each of the 3 datasets of SCLC TumorMiner. Gene expression (genexp) is expressed as log2(FPKM+1). Statistical comparisons are performed by Student t-test. **** indicates p < 0.001; ** p < 0.01; * p < 0.05. The “Download Plot Data” tab at the bottom left enables users to export the data. **B**. Snapshot of the “Mutation Variants” tab in the SCLC TumorMiner website. Number to the right indicate the percentage of samples with deleterious mutations.

**Figure S4 (related to Fig. 3).**
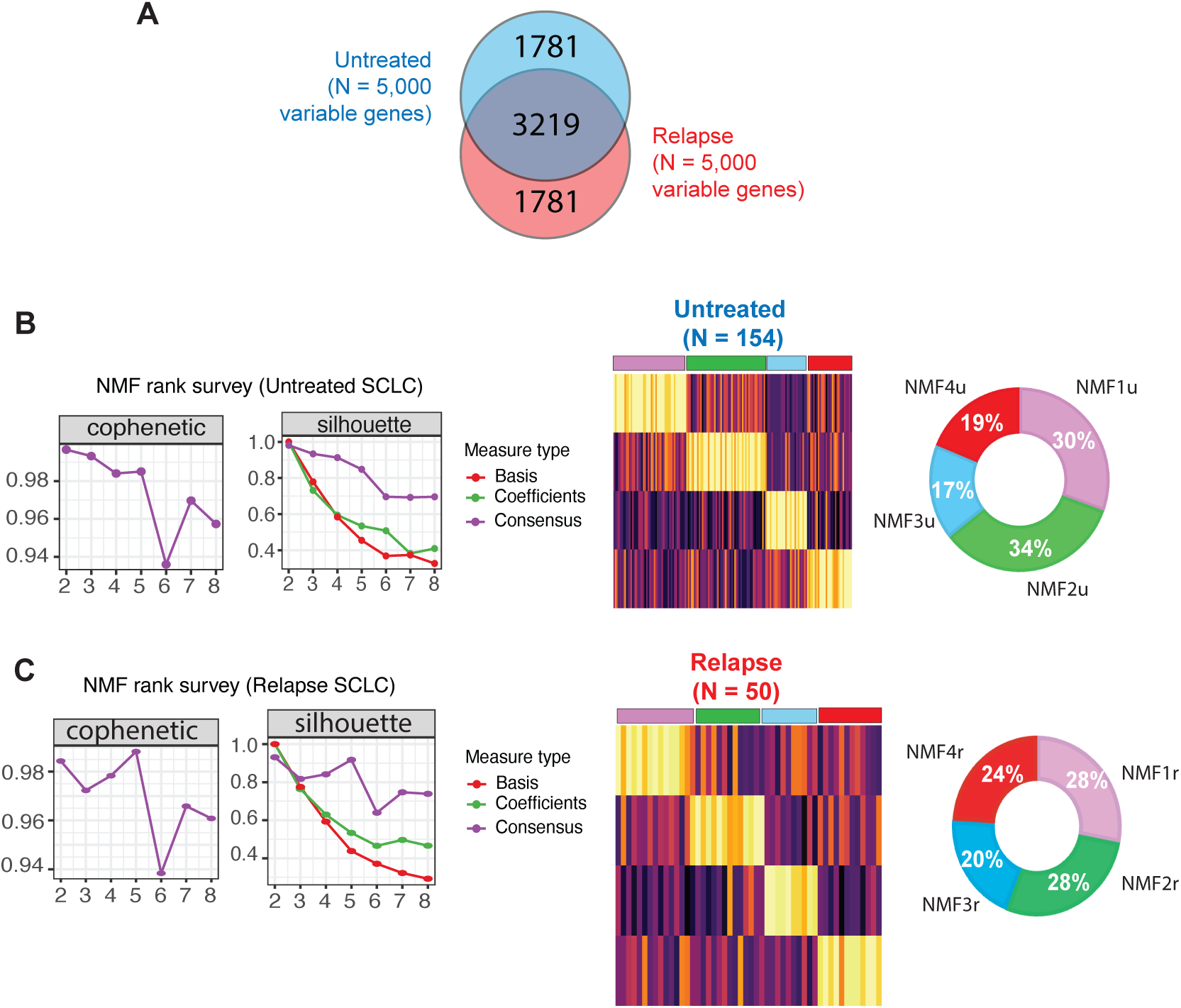
**A**. Venn-diagram showing the overlap of Untreated (top 5000) and Relapse (top 5000) most variable genes, selected using Median Absolute deviation (MAD) approach. **B** and **C** NMF rank summary for untreated and relapse SCLC samples, respectively. The cophenetic and silhouette scores are plotted for range of clusters (k=2:8), depicting k=4 as optimal number of clusters for both cohorts. Consensus matrix heatmaps at k=4 show well defined and stable clusters. Right: pie charts showing the distribution of samples in each NMF cluster.

**Figure S5 (related to Fig. 3):**
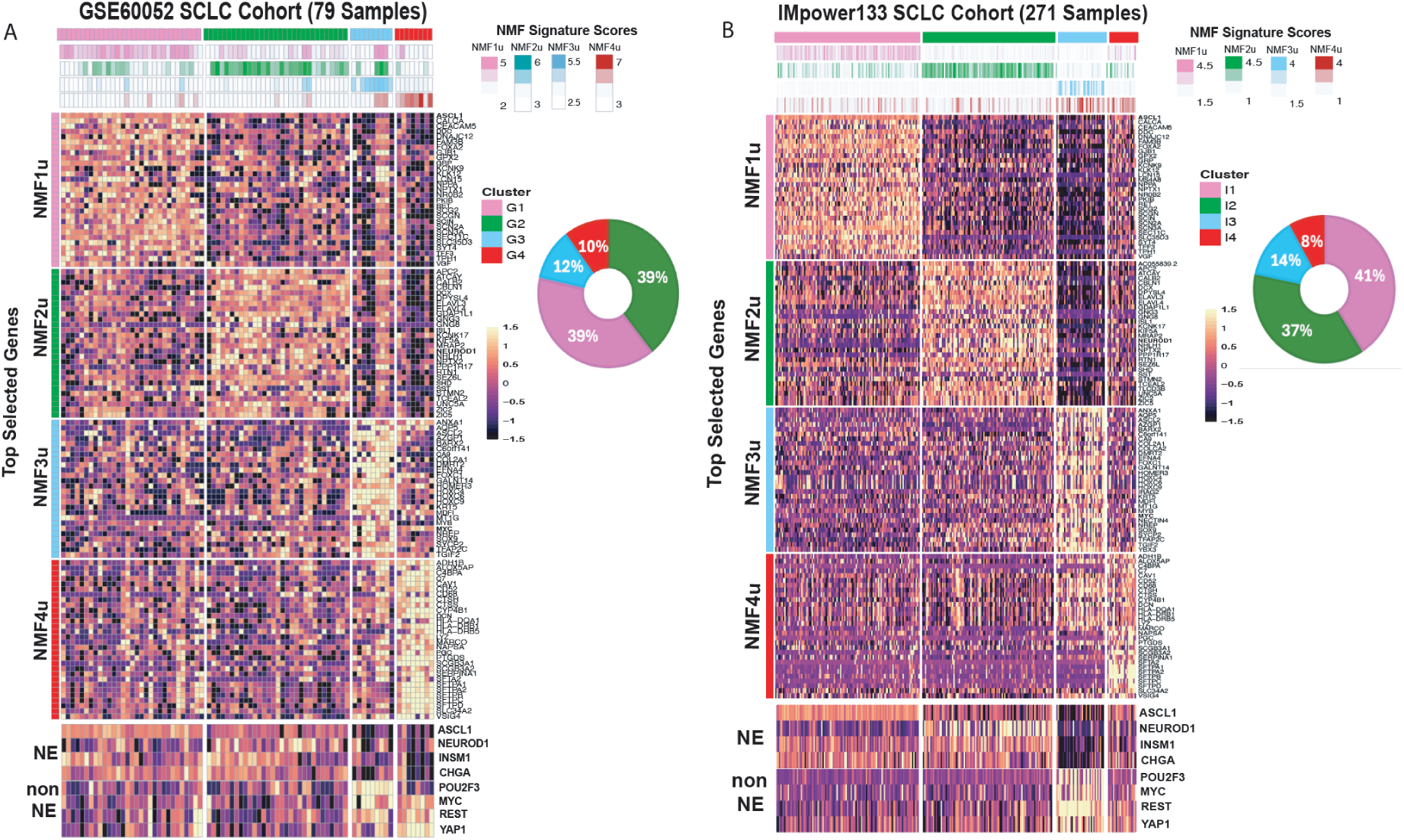
External validation of the NMFu derived gene signature on two independent SCLC cohorts (GSE60052 and Impower133). Each column represents a SCLC sample, and rows show expression of the top-30 most differentially enriched signature genes with colors indicating row-scaled expression levels. The topmost rows show 4 NMF clusters and the NMF signature scores per sample. Pie charts represent the distribution of patient samples across the different clusters.

**Figure S6 (related to Fig. 3):**
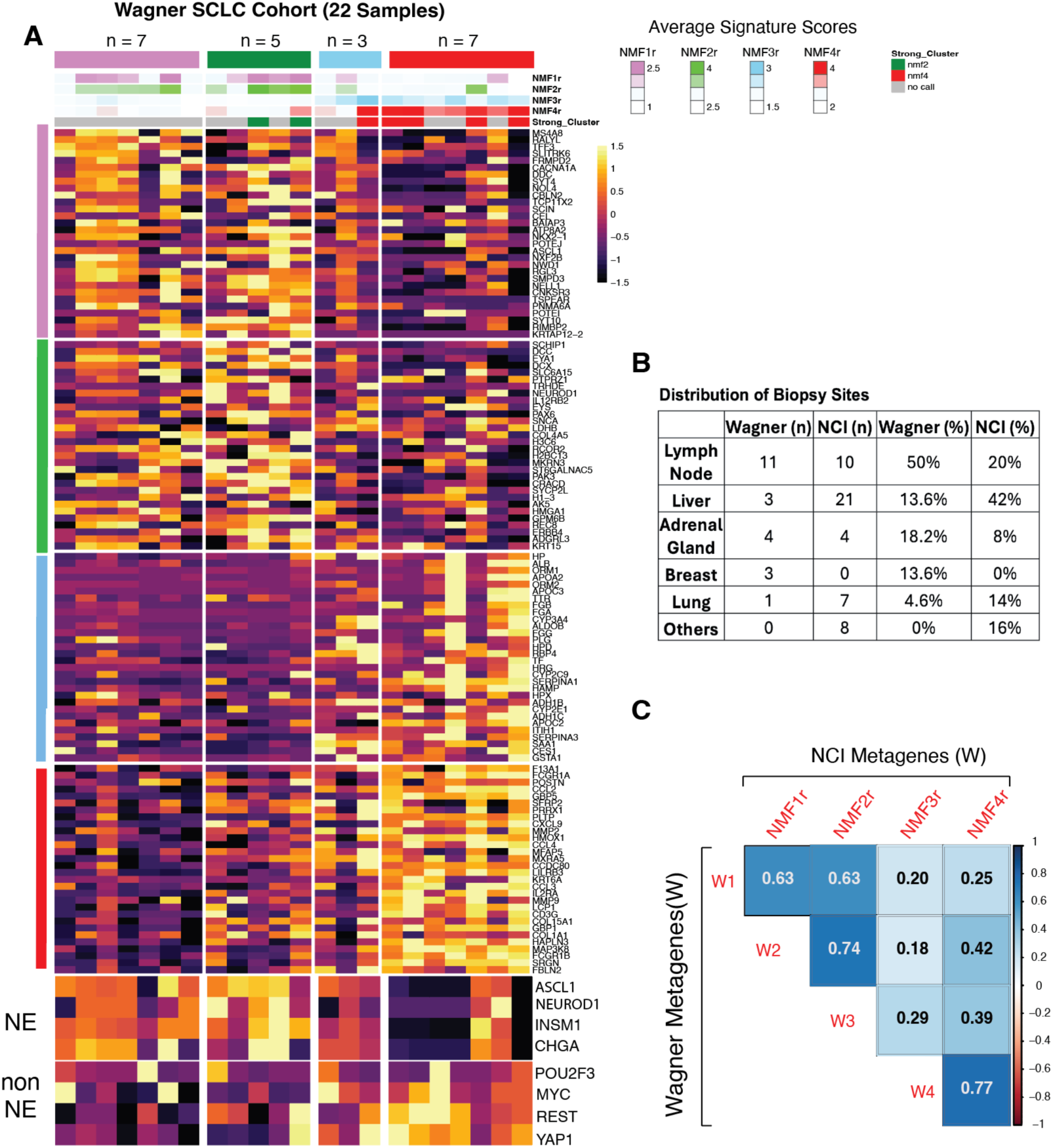
External validation of the NMFr-derived gene signature in an independent relapse SCLC cohort (Wagner et al. cohort) ^27^. **A**. Heatmap showing the expression of the top-30 most differentially enriched genes from the NCI NMFr-derived signature across the relapse samples of the Wagner et al. cohort (n = 22). Columns represent individual tumor sample, grouped by NMF cluster assignment. Rows represent the differentially enriched gene. Colors indicate row-scaled expression levels. Top annotations display NMF cluster identity and corresponding signature scores (NMFr1–NMFr4) for each sample. Bottom annotations indicate neuroendocrine (NE) and non-NE classification based on canonical markers (including *ASCL1*, *NEUROD1*, *POU2F3*, *YAP1*). **B**. Biopsy sites in the Wagner and the NCI cohorts. **C**. Concordance between the Wagner et al. cohort subgroups and NCI metagenes for each of the NMFr. Heatmap shows correlation coefficients between Wagner NMF metagene signatures (rows; W1–W4) and the NCI metagene signatures (columns; NMF1r-NMF4r), demonstrating cross-cohort reproducibility of transcriptional subtypes.

**Figure S7 (related to Fig. 7).**
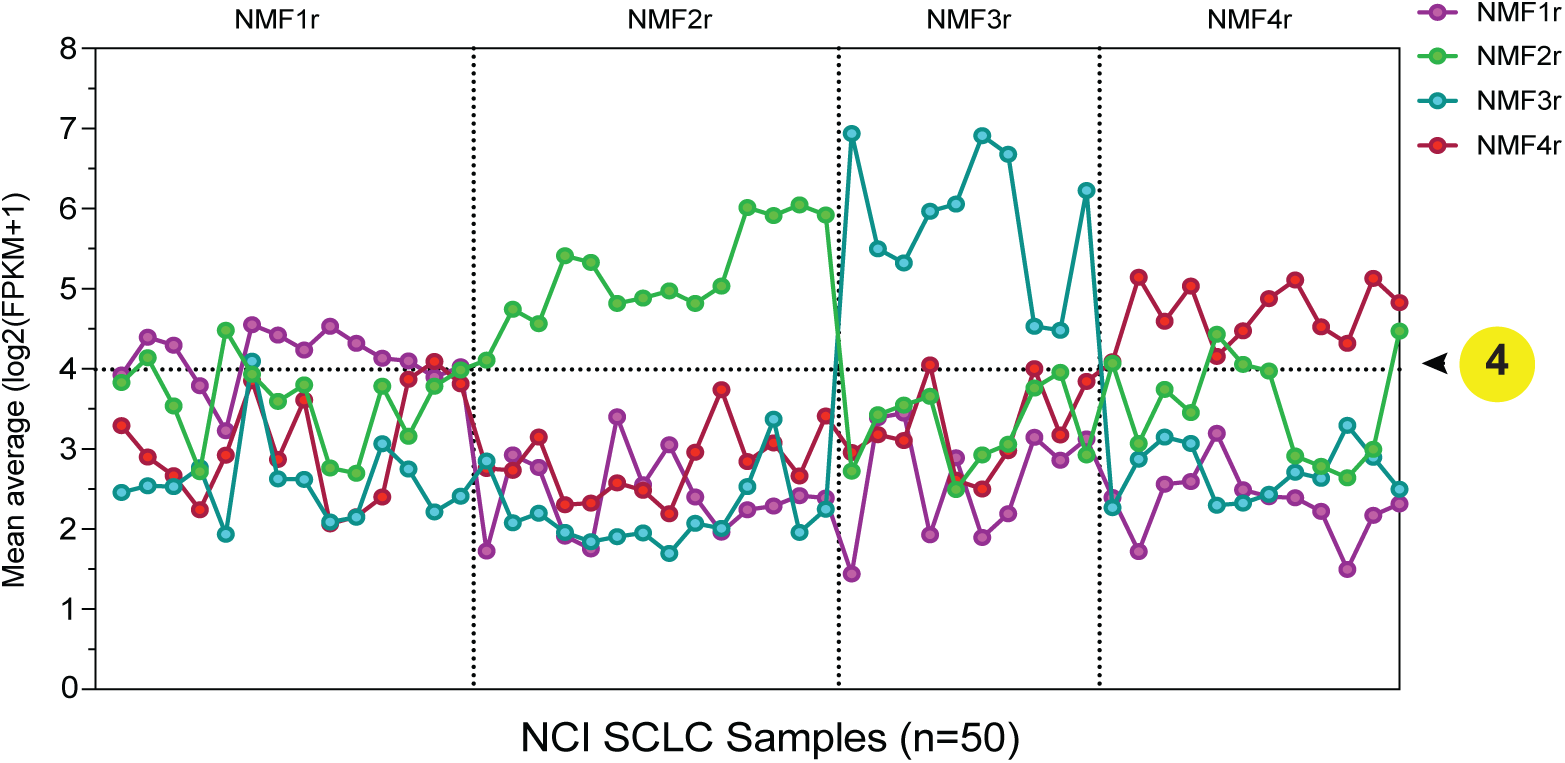
Mean average NMF expression for each and all 50 samples from patients at relapse. A mean average threshold value log2(FPKM+1) > 4 is sufficient to assign each sample to one of the 4 NMFs. The gray area defines non-significant values.

**Figure S8:**
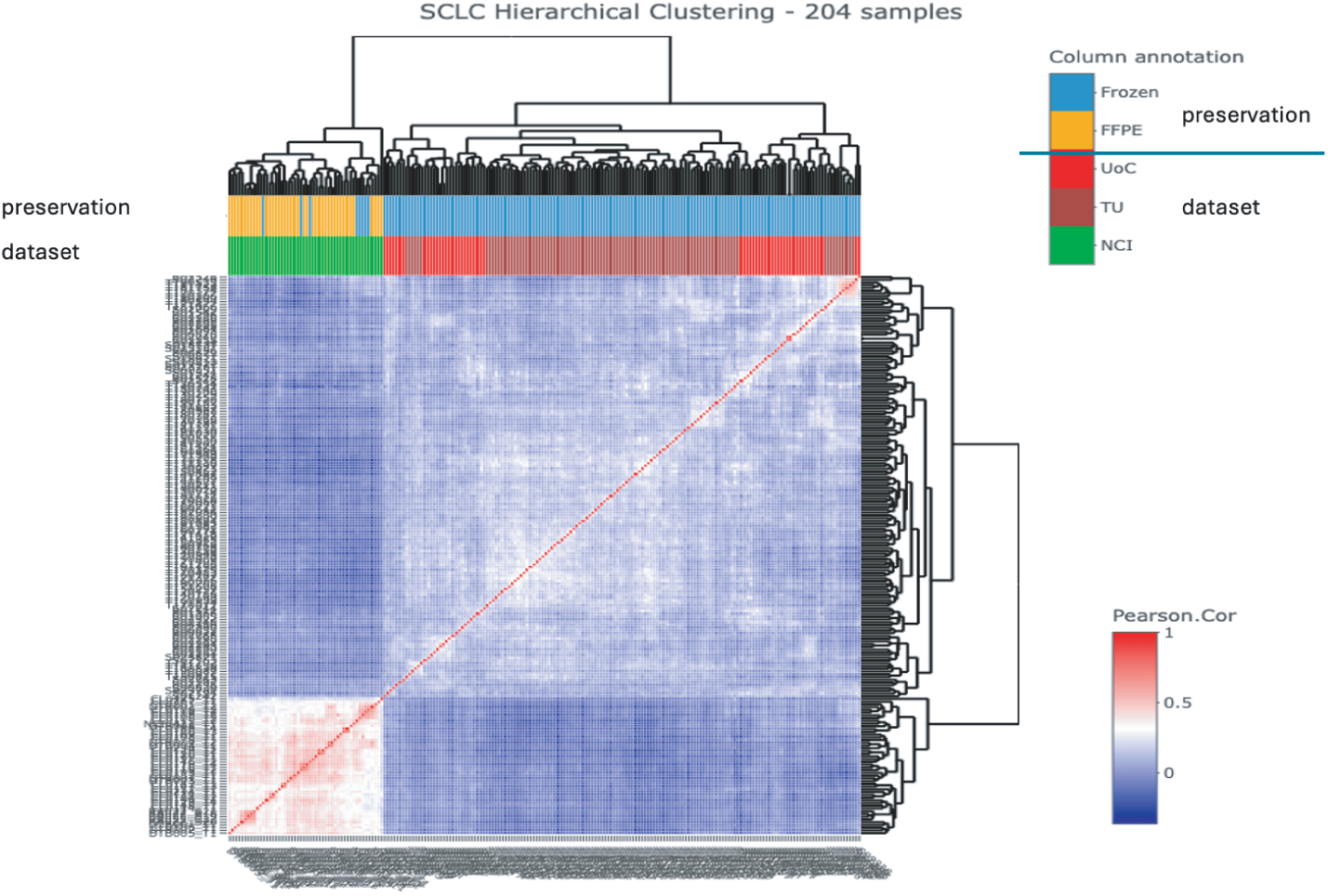
Hierarchical clustering of SCLC samples. The clustering shows a clear separation between NCI relapse samples (colored green) and untreated samples (TU, brown; UoC, red), as indicated in the first annotation row (sample type). The second annotation row (preservation method) shows that most NCI relapse samples are FFPE (orange), whereas all untreated samples are frozen (light blue). Because the preservation method is confounded with sample type (relapse vs. untreated), its effect cannot be disentangled.

## REFERENCES

1. Thomas, A., Takahashi, N., Oplustil O’Connor, L., Redon, C.E., Mohindroo, C., Sciuto, L., Pongor, L., Schmidt, K.T., Steinberg, S.M., Aladjem, M.I., et al. (2025). Tumor-targeted top1 inhibitor delivery with optimized parp inhibition in advanced solid tumors: a phase i trial of gapped scheduling. Nature Communications 16, 9457. 10.1038/s41467-025-64509-5.

2. George, J., Lim, J.S., Jang, S.J., Cun, Y., Ozretic, L., Kong, G., Leenders, F., Lu, X., Fernandez-Cuesta, L., Bosco, G., et al. (2015). Comprehensive genomic profiles of small cell lung cancer. Nature 524, 47–53. 10.1038/nature14664.

3. Rudin, C.M., Poirier, J.T., Byers, L.A., Dive, C., Dowlati, A., George, J., Heymach, J.V., Johnson, J.E., Lehman, J.M., MacPherson, D., et al. (2019). Molecular subtypes of small cell lung cancer: a synthesis of human and mouse model data. Nat Rev Cancer. 10.1038/s41568-019-0133-9.

4. Tlemsani, C., Pongor, L., Elloumi, F., Girard, L., Huffman, K.E., Roper, N., Varma, S., Luna, A., Rajapakse, V.N., Sebastian, R., et al. (2020). SCLC-CellMiner: A Resource for Small Cell Lung Cancer Cell Line Genomics and Pharmacology Based on Genomic Signatures. Cell Rep 33, 108296. 10.1016/j.celrep.2020.108296.

5. Gay, C.M., Stewart, C.A., Park, E.M., Diao, L., Groves, S.M., Heeke, S., Nabet, B.Y., Fujimoto, J., Solis, L.M., Lu, W., et al. (2021). Patterns of transcription factor programs and immune pathway activation define four major subtypes of SCLC with distinct therapeutic vulnerabilities. Cancer Cell 39, 346–360 e347. 10.1016/j.ccell.2020.12.014.

6. Pacheco, J., and Bunn, P.A. (2019). Advancements in Small-cell Lung Cancer: The Changing Landscape Following IMpower-133. Clin Lung Cancer 20, 148–160 e142. 10.1016/j.cllc.2018.12.019.

7. Paz-Ares, L., Dvorkin, M., Chen, Y., Reinmuth, N., Hotta, K., Trukhin, D., Statsenko, G., Hochmair, M.J., Ozguroglu, M., Ji, J.H., et al. (2019). Durvalumab plus platinum-etoposide versus platinum-etoposide in first-line treatment of extensive-stage small-cell lung cancer (CASPIAN): a randomised, controlled, open-label, phase 3 trial. Lancet 394, 1929–1939. 10.1016/S0140-6736(19)32222-6.

8. Nabet, B.Y., Hamidi, H., Lee, M.C., Banchereau, R., Morris, S., Adler, L., Gayevskiy, V., Elhossiny, A.M., Srivastava, M.K., Patil, N.S., et al. (2024). Immune heterogeneity in small-cell lung cancer and vulnerability to immune checkpoint blockade. Cancer Cell. 10.1016/j.ccell.2024.01.010.

9. Liu, Q., Zhang, J., Guo, C., Wang, M., Wang, C., Yan, Y., Sun, L., Wang, D., Zhang, L., Yu, H., et al. (2024). Proteogenomic characterization of small cell lung cancer identifies biological insights and subtype-specific therapeutic strategies. Cell 187, 184–203 e128. 10.1016/j.cell.2023.12.004.

10. Tang, S.W., Thomas, A., Murai, J., Trepel, J.B., Bates, S.E., Rajapakse, V.N., and Pommier, Y. (2018). Overcoming Resistance to DNA-Targeted Agents by Epigenetic Activation of Schlafen 11 (SLFN11) Expression with Class I Histone Deacetylase Inhibitors. Clin Cancer Res 24, 1944–1953. 10.1158/1078-0432.CCR-17-0443.

11. Pozo, K., Kollipara, R.K., Kelenis, D.P., Rodarte, K.E., Ullrich, M.S., Zhang, X., Minna, J.D., and Johnson, J.E. (2021). ASCL1, NKX2-1, and PROX1 co-regulate subtype-specific genes in small-cell lung cancer. iScience 24, 102953. 10.1016/j.isci.2021.102953.

12. Anderson, K.R., Torres, C.A., Solomon, K., Becker, T.C., Newgard, C.B., Wright, C.V., Hagman, J., and Sussel, L. (2009). Cooperative transcriptional regulation of the essential pancreatic islet gene NeuroD1 (beta2) by Nkx2.2 and neurogenin 3. J Biol Chem 284, 31236–31248. 10.1074/jbc.M109.048694.

13. Gosmain, Y., Marthinet, E., Cheyssac, C., Guerardel, A., Mamin, A., Katz, L.S., Bouzakri, K., and Philippe, J. (2010). Pax6 controls the expression of critical genes involved in pancreatic alpha cell differentiation and function. J Biol Chem 285, 33381–33393. 10.1074/jbc.M110.147215.

14. Mastrantonio, R., You, H., and Tamagnone, L. (2021). Semaphorins as emerging clinical biomarkers and therapeutic targets in cancer. Theranostics 11, 3262–3277. 10.7150/thno.54023.

15. Huang, Y.H., Klingbeil, O., He, X.Y., Wu, X.S., Arun, G., Lu, B., Somerville, T.D.D., Milazzo, J.P., Wilkinson, J.E., Demerdash, O.E., et al. (2018). POU2F3 is a master regulator of a tuft cell-like variant of small cell lung cancer. Genes Dev 32, 915–928. 10.1101/gad.314815.118.

16. Montoro, D.T., Haber, A.L., Biton, M., Vinarsky, V., Lin, B., Birket, S.E., Yuan, F., Chen, S., Leung, H.M., Villoria, J., et al. (2018). A revised airway epithelial hierarchy includes CFTR-expressing ionocytes. Nature 560, 319–324. 10.1038/s41586-018-0393-7.

17. Ireland, A.S., Xie, D.A., Hawgood, S.B., Barbier, M.W., Zuo, L.Y., Hanna, B.E., Lucas-Randolph, S., Tyson, D.R., Witt, B.L., Govindan, R., et al. (2025). Basal cell of origin resolves neuroendocrine-tuft lineage plasticity in cancer. Nature. 10.1038/s41586-025-09503-z.

18. Wu, X.S., He, X.Y., Ipsaro, J.J., Huang, Y.H., Preall, J.B., Ng, D., Shue, Y.T., Sage, J., Egeblad, M., Joshua-Tor, L., and Vakoc, C.R. (2022). OCA-T1 and OCA-T2 are coactivators of POU2F3 in the tuft cell lineage. Nature 607, 169–175. 10.1038/s41586-022-04842-7.

19. Abdel Wadood, N., Hollenhorst, M.I., Elhawy, M.I., Zhao, N., Englisch, C., Evers, S.B., Sabachvili, M., Maxeiner, S., Wyatt, A., Herr, C., et al. (2025). Tracheal tuft cells release ATP and link innate to adaptive immunity in pneumonia. Nature Communications 16, 584. 10.1038/s41467-025-55936-5.

20. Baine, M.K., Febres-Aldana, C.A., Chang, J.C., Jungbluth, A.A., Sethi, S., Antonescu, C.R., Travis, W.D., Hsieh, M.S., Roh, M.S., Homer, R.J., et al. (2022). POU2F3 in SCLC: Clinicopathologic and Genomic Analysis With a Focus on Its Diagnostic Utility in Neuroendocrine-Low SCLC. J Thorac Oncol 17, 1109–1121. 10.1016/j.jtho.2022.06.004.

21. Shue, Y.T., Drainas, A.P., Li, N.Y., Pearsall, S.M., Morgan, D., Sinnott-Armstrong, N., Hipkins, S.Q., Coles, G.L., Lim, J.S., Oro, A.E., et al. (2022). A conserved YAP/Notch/REST network controls the neuroendocrine cell fate in the lungs. Nat Commun 13, 2690. 10.1038/s41467-022-30416-2.

22. Kim, J.W., Ko, J.H., and Sage, J. (2022). DLL3 regulates Notch signaling in small cell lung cancer. iScience 25, 105603. 10.1016/j.isci.2022.105603.

23. Ireland, A.S., Micinski, A.M., Kastner, D.W., Guo, B., Wait, S.J., Spainhower, K.B., Conley, C.C., Chen, O.S., Guthrie, M.R., Soltero, D., et al. (2020). MYC Drives Temporal Evolution of Small Cell Lung Cancer Subtypes by Reprogramming Neuroendocrine Fate. Cancer Cell 38, 60–78 e12. 10.1016/j.ccell.2020.05.001.

24. Gazdar, A.F., Bunn, P.A., and Minna, J.D. (2017). Small-cell lung cancer: what we know, what we need to know and the path forward. Nat Rev Cancer 17, 725–737. 10.1038/nrc.2017.87.

25. Gazdar, A.F., Carney, D.N., Nau, M.M., and Minna, J.D. (1985). Characterization of variant subclasses of cell lines derived from small cell lung cancer having distinctive biochemical, morphological, and growth properties. Cancer Res 45, 2924–2930.

26. Jiang, L., Huang, J., Higgs, B.W., Hu, Z., Xiao, Z., Yao, X., Conley, S., Zhong, H., Liu, Z., Brohawn, P., et al. (2016). Genomic Landscape Survey Identifies SRSF1 as a Key Oncodriver in Small Cell Lung Cancer. PLoS Genet 12, e1005895. 10.1371/journal.pgen.1005895.

27. Wagner, A.H., Devarakonda, S., Skidmore, Z.L., Krysiak, K., Ramu, A., Trani, L., Kunisaki, J., Masood, A., Waqar, S.N., Spies, N.C., et al. (2018). Recurrent WNT pathway alterations are frequent in relapsed small cell lung cancer. Nat Commun 9, 3787. 10.1038/s41467-018-06162-9.

28. Jo, U., and Pommier, Y. (2022). Structural, molecular, and functional insights into Schlafen proteins. Experimental & Molecular Medicine. 10.1038/s12276-022-00794-0.

29. Murai, J., Thomas, A., Miettinen, M., and Pommier, Y. (2019). Schlafen 11 (SLFN11), a restriction factor for replicative stress induced by DNA-targeting anti-cancer therapies. Pharmacol Ther 201, 94–102. 10.1016/j.pharmthera.2019.05.009.

30. Masuda, K., Yoshida, T., Motoi, N., Shinno, Y., Matsumoto, Y., Okuma, Y., Goto, Y., Horinouchi, H., Yamamoto, N., Watanabe, S.-i., et al. (2025). Schlafen 11 Expression in Patients With Small Cell Lung Cancer and Its Association With Clinical Outcomes. Thoracic Cancer 16, e15529. 10.1111/1759-7714.15529.

31. Dowlati, A., Chiang, A.C., Cervantes, A., Babu, S., Hamilton, E., Wong, S.F., Tazbirkova, A., Sullivan, I.G., van Marcke, C., Italiano, A., et al. (2025). Phase 2 Open-Label Study of Sacituzumab Govitecan as Second-Line Therapy in Patients With Extensive-Stage SCLC: Results From TROPiCS-03. J Thorac Oncol. 10.1016/j.jtho.2024.12.028.

32. Ahn, M.-J., Cho, B.C., Felip, E., Korantzis, I., Ohashi, K., Majem, M., Juan-Vidal, O., Handzhiev, S., Izumi, H., Lee, J.-S., et al. (2023). Tarlatamab for Patients with Previously Treated Small-Cell Lung Cancer. New England Journal of Medicine 389, 2063–2075. 10.1056/NEJMoa2307980.

33. Sun, N.Y., Kumar, S., Kim, Y.S., Varghese, D., Mendoza, A., Nguyen, R., Okada, R., Reilly, K., Widemann, B., Pommier, Y., et al. (2025). Identification of DLK1, a Notch ligand, as an immunotherapeutic target and regulator of tumor cell plasticity and chemoresistance in adrenocortical carcinoma. Nat Commun 16, 5511. 10.1101/2024.10.09.617077.

34. Han, K., Chen, Y., Sun, X., Wen, L., Wu, Y., Chen, S., Wei, L., Yu, J., Zeng, T., Jiang, L., and Tan, L. (2025). Combining serum CDK1 with tumor markers for the diagnosis of small cell lung cancer. Clin Transl Oncol 27, 2005–2013. 10.1007/s12094-024-03722-y.

35. Muley, T., Herth, F.J., Heussel, C.P., Kriegsmann, M., Thomas, M., Meister, M., Schneider, M.A., Wehnl, B., Mang, A., and Holdenrieder, S. (2024). Prognostic value of tumor markers ProGRP, NSE and CYFRA 21-1 in patients with small cell lung cancer and chemotherapy-induced remission. Tumour Biol 46, S219–S232. 10.3233/TUB-230016.

36. Kudoh, S., Tenjin, Y., Kameyama, H., Ichimura, T., Yamada, T., Matsuo, A., Kudo, N., Sato, Y., and Ito, T. (2020). Significance of achaete-scute complex homologue 1 (ASCL1) in pulmonary neuroendocrine carcinomas; RNA sequence analyses using small cell lung cancer cells and Ascl1-induced pulmonary neuroendocrine carcinoma cells. Histochem Cell Biol 153, 443–456. 10.1007/s00418-020-01863-z.

37. Ajay, A., Wang, H., Rezvani, A., Savari, O., Grubb, B.J., McColl, K.S., Yoon, S., Joseph, P.L., Kopp, S.R., Kresak, A.M., et al. (2025). Assessment of targets of antibody drug conjugates in SCLC. NPJ Precis Oncol 9, 1. 10.1038/s41698-024-00784-7.

38. Qu, S., Fetsch, P., Thomas, A., Pommier, Y., Schrump, D.S., Miettinen, M.M., and Chen, H. (2022). Molecular Subtypes of Primary SCLC Tumors and Their Associations With Neuroendocrine and Therapeutic Markers. J Thorac Oncol 17, 141–153. 10.1016/j.jtho.2021.08.763.

39. Sun, F.D., and Das, M.S. (2024). Targeting the EZH2-SLFN11 pathway-a lesson in developing molecularly-informed treatments for recurrent small cell lung cancer. Translational Cancer Research 13, 6608–6612. 10.21037/tcr-24-1755.

40. Zhang, B., Stewart, C.A., Wang, Q., Cardnell, R.J., Rocha, P., Fujimoto, J., Solis Soto, L.M., Wang, R., Novegil, V., Ansell, P., et al. (2022). Dynamic expression of Schlafen 11 (SLFN11) in circulating tumour cells as a liquid biomarker in small cell lung cancer. Br J Cancer. 10.1038/s41416-022-01811-9.

41. Kundu, K., Cardnell, R.J., Zhang, B., Shen, L., Stewart, C.A., Ramkumar, K., Cargill, K.R., Wang, J., Gay, C.M., and Byers, L.A. (2021). SLFN11 biomarker status predicts response to lurbinectedin as a single agent and in combination with ATR inhibition in small cell lung cancer. Transl Lung Cancer Res 10, 4095–4105. 10.21037/tlcr-21-437.

42. Laroche-Clary, A., Chaire, V., Valverde, V., Derieppe, M.A., and Italiano, A. (2025). Synergistic Effects of ATR Inhibition and Lurbinectedin in Soft-Tissue Sarcomas: The Predictive Role of SLFN11 Expression. Clin Cancer Res. 10.1158/1078-0432.CCR-24-2556.

43. Thomas, A., Tanaka, M., Trepel, J., Reinhold, W.C., Rajapakse, V.N., and Pommier, Y. (2017). Temozolomide in the Era of Precision Medicine. Cancer Res 77, 823–826. 10.1158/0008-5472.CAN-16-2983.

44. Schultz, C.W., Zhang, Y., Elmeskini, R., Zimmermann, A., Fu, H., Murai, Y., Wangsa, D., Kumar, S., Takahashi, N., Atkinson, D., et al. (2023). ATR inhibition augments the efficacy of lurbinectedin in small-cell lung cancer. EMBO Molecular Medicine 15, e17313. 10.15252/emmm.202217313.

45. Gao, J., Aksoy, B.A., Dogrusoz, U., Dresdner, G., Gross, B., Sumer, S.O., Sun, Y., Jacobsen, A., Sinha, R., Larsson, E., et al. (2013). Integrative analysis of complex cancer genomics and clinical profiles using the cBioPortal. Sci Signal 6, pl1. 10.1126/scisignal.2004088.

46. Goldman, M.J., Craft, B., Hastie, M., Repečka, K., McDade, F., Kamath, A., Banerjee, A., Luo, Y., Rogers, D., Brooks, A.N., et al. (2020). Visualizing and interpreting cancer genomics data via the Xena platform. Nat Biotechnol 38, 675–678. 10.1038/s41587-020-0546-8.

47. Tang, Z., Kang, B., Li, C., Chen, T., and Zhang, Z. (2019). GEPIA2: an enhanced web server for large-scale expression profiling and interactive analysis. Nucleic Acids Res 47, W556–W560. 10.1093/nar/gkz430.

48. Chandrashekar, D.S., Karthikeyan, S.K., Korla, P.K., Patel, H., Shovon, A.R., Athar, M., Netto, G.J., Qin, Z.S., Kumar, S., Manne, U., et al. (2022). UALCAN: An update to the integrated cancer data analysis platform. Neoplasia 25, 18–27. 10.1016/j.neo.2022.01.001.

49. Polley, E., Kunkel, M., Evans, D., Silvers, T., Delosh, R., Laudeman, J., Ogle, C., Reinhart, R., Selby, M., Connelly, J., et al. (2016). Small Cell Lung Cancer Screen of Oncology Drugs, Investigational Agents, and Gene and microRNA Expression. J Natl Cancer Inst 108. 10.1093/jnci/djw122.

50. Simpson, K.L., Rothwell, D.G., Blackhall, F., and Dive, C. (2025). Challenges of small cell lung cancer heterogeneity and phenotypic plasticity. Nat Rev Cancer 25, 447–462. 10.1038/s41568-025-00803-0.

51. Chan, J.M., Quintanal-Villalonga, Á., Gao, V.R., Xie, Y., Allaj, V., Chaudhary, O., Masilionis, I., Egger, J., Chow, A., Walle, T., et al. (2021). Signatures of plasticity, metastasis, and immunosuppression in an atlas of human small cell lung cancer. Cancer Cell 39, 1479–1496.e1418. 10.1016/j.ccell.2021.09.008.

52. Lissa, D., Takahashi, N., Desai, P., Manukyan, I., Schultz, C.W., Rajapakse, V., Velez, M.J., Mulford, D., Roper, N., Nichols, S., et al. (2022). Heterogeneity of neuroendocrine transcriptional states in metastatic small cell lung cancers and patient-derived models. Nat Commun 13, 2023. 10.1038/s41467-022-29517-9.

53. Stewart, C.A., Gay, C.M., Xi, Y., Sivajothi, S., Sivakamasundari, V., Fujimoto, J., Bolisetty, M., Hartsfield, P.M., Balasubramaniyan, V., Chalishazar, M.D., et al. (2020). Single-cell analyses reveal increased intratumoral heterogeneity after the onset of therapy resistance in small-cell lung cancer. Nature Cancer. 10.1038/s43018-019-0020-z.

54. Zhang, W., Girard, L., Zhang, Y.A., Haruki, T., Papari-Zareei, M., Stastny, V., Ghayee, H.K., Pacak, K., Oliver, T.G., Minna, J.D., and Gazdar, A.F. (2018). Small cell lung cancer tumors and preclinical models display heterogeneity of neuroendocrine phenotypes. Transl Lung Cancer Res 7, 32–49. 10.21037/tlcr.2018.02.02.

55. Ireland, A.S., Hawgood, S.B., Xie, D.A., Barbier, M.W., Lucas-Randolph, S., Tyson, D.R., Zuo, L.Y., Witt, B.L., Govindan, R., Dowlati, A., et al. (2024). Basal cell of origin resolves neuroendocrine–tuft lineage plasticity in cancer. bioRxiv, 2024.2011.2013.623500. 10.1101/2024.11.13.623500.

56. Qi, J., Zhang, J., Liu, N., Zhao, L., and Xu, B. (2022). Prognostic Implications of Molecular Subtypes in Primary Small Cell Lung Cancer and Their Correlation With Cancer Immunity. Front Oncol 12, 779276. 10.3389/fonc.2022.779276.

57. Borromeo, M.D., Savage, T.K., Kollipara, R.K., He, M., Augustyn, A., Osborne, J.K., Girard, L., Minna, J.D., Gazdar, A.F., Cobb, M.H., and Johnson, J.E. (2016). ASCL1 and NEUROD1 Reveal Heterogeneity in Pulmonary Neuroendocrine Tumors and Regulate Distinct Genetic Programs. Cell Rep 16, 1259–1272. 10.1016/j.celrep.2016.06.081.

58. Choudhury, N.J., Lai, W.V., Makhnin, A., Heller, G., Eng, J., Li, B., Preeshagul, I., Santini, F.C., Offin, M., Ng, K., et al. (2024). A Phase I/II Study of Valemetostat (DS-3201b), an EZH1/2 Inhibitor, in Combination with Irinotecan in Patients with Recurrent Small-Cell Lung Cancer. Clin Cancer Res 30, 3697–3703. 10.1158/1078-0432.CCR-23-3383.

59. Rudin, C.M., Reck, M., Johnson, M.L., Blackhall, F., Hann, C.L., Yang, J.C., Bailis, J.M., Bebb, G., Goldrick, A., Umejiego, J., and Paz-Ares, L. (2023). Emerging therapies targeting the delta-like ligand 3 (DLL3) in small cell lung cancer. J Hematol Oncol 16, 66. 10.1186/s13045-023-01464-y.

60. Paul, S., Konig, M.F., Pardoll, D.M., Bettegowda, C., Papadopoulos, N., Wright, K.M., Gabelli, S.B., Ho, M., van Elsas, A., and Zhou, S. (2024). Cancer therapy with antibodies. Nat Rev Cancer 24, 399–426. 10.1038/s41568-024-00690-x.

61. Shen, L., Sun, X., Chen, Z., Guo, Y., Shen, Z., Song, Y., Xin, W., Ding, H., Ma, X., Xu, W., et al. (2024). ADCdb: the database of antibody-drug conjugates. Nucleic Acids Res 52, D1097–D1109. 10.1093/nar/gkad831.

62. DeLucia, D.C., Cardillo, T.M., Ang, L., Labrecque, M.P., Zhang, A., Hopkins, J.E., De Sarkar, N., Coleman, I., da Costa, R.M.G., Corey, E., et al. (2021). Regulation of CEACAM5 and Therapeutic Efficacy of an Anti-CEACAM5-SN38 Antibody-drug Conjugate in Neuroendocrine Prostate Cancer. Clin Cancer Res 27, 759–774. 10.1158/1078-0432.CCR-20-3396.

63. Du, Y., Wen, Y., and Huang, J. (2024). Analysis of variation of serum CEA, SCC, CYFRA21-1 in patients with lung cancer and their diagnostic value with EBUS-TBNA. J Med Biochem 43, 363–371. 10.5937/jomb0-37083.

64. Yu, M., and Wang, P. (2024). A comparative study of ProGRP and CEA as serological markers in small cell lung cancer treatment. Discov Oncol 15, 485. 10.1007/s12672-024-01323-3.

65. Kim, Y.S., Nabet, B.Y., Cortez, B.N., Sun, N.Y., Sebastian, R., Redon, C.E., Ray, A., Liu, L., Ishola, A.A., Loew, S., et al. (2025). NOTCH1 reverses immune suppression in small cell lung cancer through reactivation of STING. J Clin Invest 135. 10.1172/JCI185423.

66. Roper, N., Velez, M.J., Chiappori, A., Kim, Y.S., Wei, J.S., Sindiri, S., Takahashi, N., Mulford, D., Kumar, S., Ylaya, K., et al. (2021). Notch signaling and efficacy of PD-1/PD-L1 blockade in relapsed small cell lung cancer. Nat Commun 12, 3880. 10.1038/s41467-021-24164-y.

67. Mayakonda, A., Lin, D.C., Assenov, Y., Plass, C., and Koeffler, H.P. (2018). Maftools: efficient and comprehensive analysis of somatic variants in cancer. Genome Res 28, 1747–1756. 10.1101/gr.239244.118.

